# Single-nucleus and spatial transcriptomics of archival pancreatic cancer reveals multi-compartment reprogramming after neoadjuvant treatment

**DOI:** 10.1101/2020.08.25.267336

**Authors:** William L. Hwang, Karthik A. Jagadeesh, Jimmy A. Guo, Hannah I. Hoffman, Payman Yadollahpour, Rahul Mohan, Eugene Drokhlyansky, Nicholas Van Wittenberghe, Orr Ashenberg, Samouil Farhi, Denis Schapiro, Jason Reeves, Daniel R. Zollinger, George Eng, Jason M. Schenkel, William A. Freed-Pastor, Clifton Rodrigues, Joshua Gould, Conner Lambden, Caroline Porter, Alexander Tsankov, Danielle Dionne, Domenic Abbondanza, Julia Waldman, Michael Cuoco, Lan Nguyen, Toni Delorey, Devan Phillips, Debora Ciprani, Marina Kern, Arnav Mehta, Kit Fuhrman, Robin Fropf, Joseph Beechem, Jay S. Loeffler, David P. Ryan, Colin D. Weekes, David T. Ting, Cristina R. Ferrone, Jennifer Y. Wo, Theodore S. Hong, Andrew J. Aguirre, Orit Rozenblatt-Rosen, Mari Mino-Kenudson, Carlos Fernandez-del Castillo, Andrew S. Liss, Tyler Jacks, Aviv Regev

**Author notes:** These authors contributed equally: William L. Hwang, Karthik A. Jagadeesh. These authors jointly supervised this work: Aviv Regev, Tyler E. Jacks. Genentech, 1 DNA Way, South San Francisco, CA, USA.

## Abstract

Pancreatic ductal adenocarcinoma (PDAC) remains a treatment-refractory disease. Characterizing PDAC by mRNA profiling remains particularly challenging. Previously identified bulk expression subtypes were influenced by contaminating stroma and have not yet informed clinical management, whereas single cell RNA-seq (scRNA-seq) of fresh tumors under-represented key cell types. Here, we developed a robust single-nucleus RNA-seq (snRNA-seq) technique for frozen archival PDAC specimens and used it to study both untreated tumors and those that received neoadjuvant chemotherapy and radiotherapy (CRT). Gene expression programs learned across untreated malignant cell and fibroblast profiles uncovered a clinically relevant molecular taxonomy with improved prognostic stratification compared to prior classifications. Moreover, in the increasingly-adopted neoadjuvant treatment context, there was a depletion of classical-like phenotypes in malignant cells in favor of basal-like phenotypes associated with TNF-NFkB and interferon signaling as well as the presence of novel acinar and neuroendocrine classical-like states, which may be more resilient to cytotoxic treatment. Spatially-resolved transcriptomics revealed an association between malignant cells expressing these basal-like programs and higher immune infiltration with increased lymphocytic content, whereas those exhibiting classical-like programs were linked to sparser macrophage-predominant microniches, perhaps pointing to susceptibility to distinct therapeutic strategies. Our refined molecular taxonomy and spatial resolution can help advance precision oncology in PDAC through informative stratification in clinical trials and insights into differential therapeutic targeting leveraging the immune system.

## INTRODUCTION

Pancreatic ductal adenocarcinoma (PDAC) is projected to become the second leading cause of cancer death in the United States by 2030 (*1, 2*). Despite advancements in systemic therapy, many patients cannot receive post-operative chemotherapy and/or radiotherapy (CRT) due to the morbidity often associated with surgery (*3, 4*). Neoadjuvant therapy has been increasingly adopted to aggressively address the risk of micrometastatic spread and to circumvent concerns of treatment tolerance in the postoperative setting. Recent clinical trials support the use of preoperative treatment over upfront surgery (*5, 6*). Nonetheless, tumor resistance to CRT remains a profound challenge for PDAC. In particular, there is an urgent need to understand how preoperative treatment impacts residual tumor cells to identify additional therapeutic vulnerabilities that can be exploited in combination with neoadjuvant CRT.

Unlike many other common cancers, molecular subtyping of pancreatic cancer remains in its nascent stages and does not currently inform clinical management or therapeutic development (*7*). Bulk RNA profiling of PDAC (*8*–*13*) identified two consensus subtypes: (1) *classical-pancreatic*, encompassing a spectrum of pancreatic lineage precursors, and (2) *basal-like/squamous/quasi-mesenchymal*, characterized by loss of endodermal identity and aberrations in chromatin modifiers (*7*). Basal-like tumors were associated with worse survival and poorer responses to chemotherapy in the metastatic setting (*14*), but attempts to refine this classification (beyond basal-like *vs*. classical) have failed to further stratify patient survival (*7*). Additional identified subtypes, such as exocrine, aberrantly-differentiated endocrine exocrine (ADEX), and immunogenic, are associated with lower tumor purity and may represent stromal contributions rather than neoplastic cell-intrinsic programs (*8, 10, 11*). Bulk profiling studies attempted to address this challenge by enriching for neoplastic content by either specimen selection or microdissection (*8*–*11*), but these introduced bias and precluded the assignment of transcripts to specific cell types. Thus, it remains unclear if genes expressed in ‘normal’ differentiated pancreatic tissue have a role in cancer or represent non-malignant tissue contamination. Furthermore, most studies have been performed in the untreated setting and do not offer insights into optimal treatment approaches after CRT.

The importance of distinguishing the relationships among malignant, stromal and immune cells, is further emphasized by how the therapeutic effect of cytotoxic treatments may be mediated through their impact on the immune system (*15, 16*) or cancer-associated fibroblasts (CAFs). Specifically, CRT can stimulate the production of type I interferons (IFN), which favor a cDC1-polarized phenotype with improved capacity for cross-presentation (*17*–*22*). However, interferon signaling can also hinder anti-tumor responses in a dose- and cancer-type-dependent manner (*23, 24*), by diminishing antigen processing and upregulating nonclassical MHC class I molecules that suppress NK- and T-cell killing (*23*). These effects can negatively impact native anti-tumor immunity and immunotherapy (*23*). Activated CAFs in PDAC can also contribute to tumor growth, therapeutic resistance and immune cell exclusion (*25*–*27*). This has led to therapeutic interest in inhibiting certain CAF subpopulations, such as myofibroblasts, though surprisingly, deletion of αSMA^+^ CAFs correlated with reduced survival in transgenic mice (*28*). Such studies, however, were performed in the untreated context, and it remains unclear whether such observations would persist after neoadjuvant CRT.

Single cell RNA-seq (scRNA-seq) can help tackle these questions by distinguishing the diversity of malignant and non-malignant cells in the tumor (*29*–*32*), and elucidating the impact of therapy on each compartment and their interactions. However, scRNA-seq in PDAC has lagged behind other cancer types due to high intrinsic nuclease content and dense desmoplastic stroma (*33*–*36*), resulting in reduced RNA quality, low numbers of viable cells, preferential capture of certain cell types at the expense of others, and challenges with dissociating treated tumors. Single nucleus RNA-seq (snRNA-seq) provides a compelling alternative for difficult-to-dissociate specimens or frozen archival samples (*37*–*40*), and can better recover malignant cells and stroma while reducing stress signatures (introduced by dissociation) and maintaining the same spectrum of cell states (*41*–*43*).

Here, we optimized snRNA-seq for frozen archival PDAC specimens, largely circumventing the challenges that have hampered scRNA-seq in PDAC. We successfully applied snRNA-seq to tumors from untreated patients and from those who received neoadjuvant CRT prior to resection(*5*), including archival samples stored for up to seven years before processing, with comparable quality for untreated and treated tumors. The recovered cellular composition closely matched the tumor composition as determined by a gold standard of multiplex protein profiling *in situ*. This allowed us to discover treatment-associated remodeling of the malignant, fibroblast, and immune compartments; identify changes in expression programs in malignant cells and fibroblasts associated with treatment; and refine the molecular taxonomy of PDAC in a clinically relevant manner. Combining these expression programs with spatially-resolved transcriptomics, we associated neoplastic- and fibroblast-intrinsic programs with different local immune microenvironments, highlighting how immunomodulatory strategies may be better selected and deployed. Our work provides a window into treatment selection pressures and the changes they may induce in the molecular composition of tumors and provides a blueprint for exploring therapeutic strategies tailored for the reprogrammed tumor.

## RESULTS

### Single-nucleus RNA-seq Accurately Represents the Malignant and Non-Malignant Compartments of Human PDAC Tumors

We performed snRNA-seq on flash frozen, histologically-confirmed, primary PDAC specimens from patients (*n* = 26) with resectable or borderline-resectable disease, who underwent surgical resection with (*n* = 11) or without (*n* = 15) neoadjuvant CRT (**Figure 1A; Table S1**), and analyzed 138,547 high quality single nucleus profiles (**Methods**). We separately grouped single nucleus profiles by treatment status from all patients by unsupervised clustering and annotated cell subsets using known cell type-specific gene signatures (**Figures 1B, S1; Methods**). The identity of malignant cells was confirmed by inferred Copy Number Aberrations (CNAs) (**Figure S2A**) (*29*). The frequencies of inferred chromosome arm-level somatic CNAs in the malignant cells of these patient samples were comparable to those in The Cancer Genome Atlas (TCGA) pancreatic adenocarcinoma cohort as assessed by single nucleotide polymorphisms and whole-exome sequencing (**Figure S2B**) (*11*). Among non-malignant cells, we identified all major cell types known to compose exocrine pancreatic tumors (**Figures 1B-C**). We noted the presence of a subset of atypical ductal-like cells (*CFTR*^high^;*KRT*19^high^;CNA^low^) (**Figure 1B**) that are unlikely to be doublets based on their typical number of unique molecular identifiers, and could be further explored in future studies as possible precursors to invasive cancer cells.

**Figure 1.**
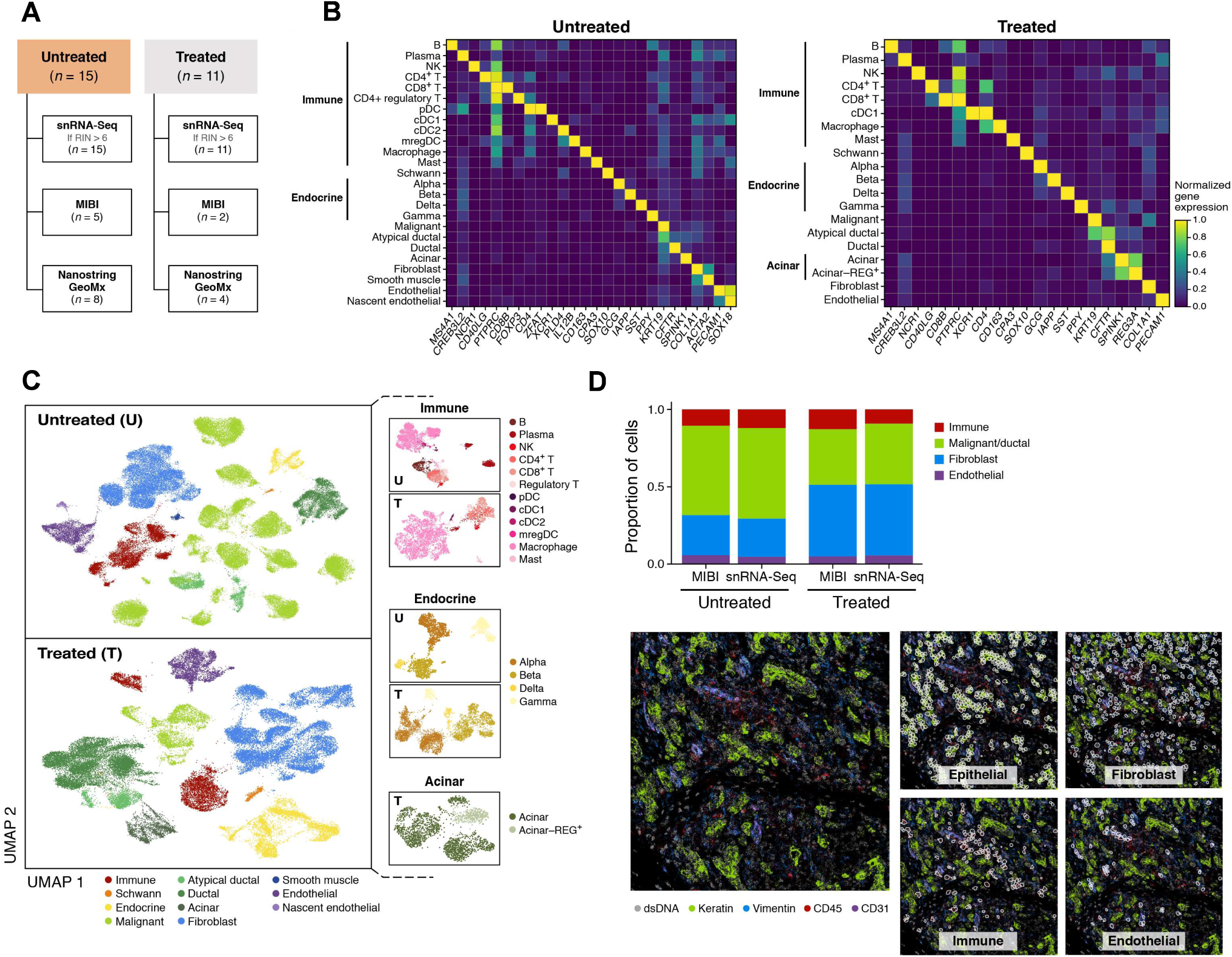
Single-nucleus RNA-seq of PDAC Captures Representative Cell Type Distributions across Malignant, Epithelial, Immune and Stromal Compartments. (A) Experimental workflow of human PDAC tumors for snRNA-seq, Multiplex Ion Beam Imaging (MIBI), and/or digital spatial profiling (NanoString GeoMx). (B,C) snRNA-seq captures diverse malignant, epithelial, immune and stromal cell subsets. (B) Mean expression (color bar) selected marker genes (columns) across annotated cell subsets (rows) of different compartments (labels, left) from untreated (left) and treated (right) tumors. (C) UMAP embedding of single nucleus profiles (dots) from untreated (top) and treated (bottom) tumors colored by *post hoc* cell type annotations (color legend). Insets: UMAP re-embedding of single nucleus profiles from specific subsets of interest. (D) snRNA-seq captures representative cell types distributions compared to *in situ* assessment. Top: Proportion of cells (y axis) in each of the four major compartments (color legend) as estimated by snRNA-seq or MIBI (x axis) in aggregate across all untreated (left; n = 5) or treated (right, n = 2) tumors. Bottom: Representative MIBI images and segmentation showing staining with antibodies against cytokeratin (green), vimentin (blue), CD45 (red), CD31 (purple) and double-stranded DNA (gray).

Examining treatment-naïve and neoadjuvant-treated specimens separately (**Figure 1C**) (*44*), non-malignant cell subsets primarily partitioned by cell type with substantial inter-patient mixing, whereas malignant cells partitioned by patient, as we previously reported for other tumor types (*29, 31, 32, 45, 46*). Among non-malignant cells, we readily annotated diverse immune, endocrine, and acinar cells, and their cell subsets by known gene signatures (**Figures 1B-C**) (*40, 47*–*49*). While earlier scRNA-seq studies in PDAC did not fully capture the stromal milieu with marked underrepresentation of cancer-associated fibroblasts (CAFs) (*50*–*52*), they are well-represented in all our samples (**Figures 1B-C, S1**).

To further assess if our method captured representative cell type proportions, we compared it to estimates from Multiplexed Ion Beam Imaging (MIBI), using a 27-plex epithelial oncology panel on formalin-fixed paraffin-embedded (FFPE) sections derived from tumor specimens in a subset of seven individuals (**Figure 1A; Methods**) (*53, 54*). This confirmed that snRNA-seq captures a representative distribution of the major cell types present in PDAC, both in aggregate across all tumors (**Figure 1D**), and individually (**Figure S3**).

### Compartment-Specific Remodeling of Cell Composition and Intrinsic Programs following Neoadjuvant Treatment has Implications for Anti-Tumor Immunity

We compared the snRNA-seq cell type proportions for the treated and untreated cohorts (**Figures 2A, S4**). As expected, the proportion of malignant cells was significantly lower in the treated cohort, which we confirmed by histology (p < 5 × 10^−5^, Fisher’s exact test). In contrast, within the non-malignant compartment, there were proportionally more acinar cells (p < 5 × 10^−5^, Fisher’s exact test), endocrine cells (p < 5 × 10^−5^), and Schwann cells (p < 5 × 10^−5^) associated with neoadjuvant CRT (**Figures 2A, S4**). This was consistent with the higher density of regenerating pancreatic tissue and the previously described resistance of some of these cell types to cytotoxic therapy (*55*). Interestingly, after removal of malignant cells, there were proportionally fewer fibroblasts in the CRT cohort (relative to immune and other cells) compared to the untreated group (**Figure S4**), suggesting that the histologically-apparent enhanced desmoplastic reaction after treatment may not be dependent on CAF proliferation but rather an enrichment in CAF phenotypes that contribute to the desmoplastic reaction (*e*.*g*., myofibroblasts) (*56*). There were substantially more Schwann cells in the CRT cohort despite their known radiosensitivity (*57*), which could be due to an active repopulation or recruitment of Schwann cells and their associated nerves to the site of treatment-induced injury. Notably, we also detected a population of regenerating acinar cells (*REG3A*^high^;*REG3G*^high^;*SYCN*^high^) in the treated specimens (**Figure 1C**), which have been associated with acinar-to-ductal metaplasia and pancreatic intraepithelial neoplasia (*40, 48, 58, 59*).

**Figure 2.**
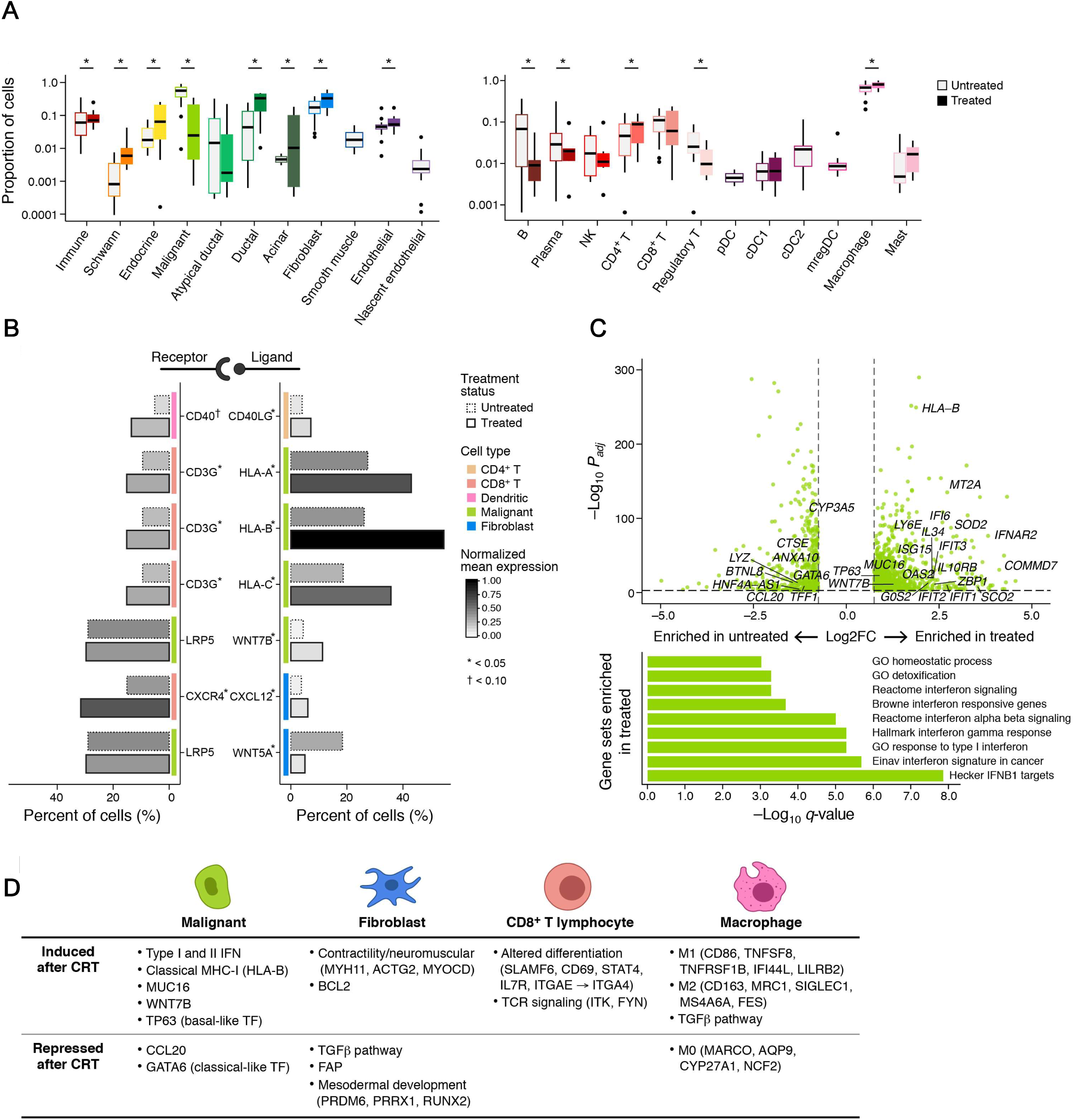
Neoadjuvant Chemoradiotherapy Remodels Cellular Subsets, Programs and Interactions in Compartment-Specific Manner. (A) CRT remodels cell type composition across compartments. Proportions (y axis) of cell subsets (x axis) in untreated (n = 15) *vs*. treated (n = 11) patients of all cells in the tumor (left) and of immune cell only (right). * p < 0.05, Fisher’s exact test. (B) Treatment impact on putative cell interactions. Selected putative interactions between cell subsets (y-axis, color code) differentially impacted by treatment status based on expression of the receptor (left) in one cell subset, and its cognate ligand (right) in another (**Methods**), distinguishing the expression level (color bar) and proportion of expressing cells (x axis) in untreated (solid borders) and treated (dashed borders) tumors. * p <0.05, † p < 0.1; Fisher’s exact test or chi-square test with Yates correction. (C) Interferon signaling and basal-like genes induced in malignant cells from CRT-treated tumors. Top: differential expression (log2(fold-change), x axis) and its significance (-log10(adjusted p-value), *y* axis, DESeq2 R package) of genes in malignant cells between treated and untreated tumors. Names of selected significant genes are marked. Bottom: Gene Set Enrichment Analysis (GSEA(*143, 149, 150*)) terms (y axis) ranked by increasing significance (-log_10_(FDR q-value)) of enrichment for induction in treated tumors. (D) Summary of compartment specific effects of CRT.

The immune compartment from treated tumors was distinct from that of the untreated tumors: there were significantly lower proportions of B cells (Fisher’s exact p < 5 × 10^−5^), plasma cells (p < 5 × 10^−5^), and regulatory T cells (p < 5 × 10^−5^) but higher proportions of CD4^+^ T cells (p = 1 × 10^−4^) and macrophages (p < 5 × 10^−5^) (**Figures 2A, S4**). We also observed a marked difference in dendritic cell (DC) subsets. First, conventional type 2 dendritic cells (cDC2), plasmacytoid dendritic cells (pDC) and mature regulatory dendritic cells (mregDCs) (*60*–*62*), which can suppress anti-tumor immunity in certain contexts, were prominent DC subtypes in treatment-naïve samples, but absent from CRT specimens (**Figure 2A**) (*63*). In contrast, conventional type 1 dendritic cells (cDC1), which activate cytotoxic lymphocytes critical for anti-tumor immunity, was the only DC subset detected in post-CRT tumors (**Figure 2A**). Second, receptor-ligand analysis inferred enhanced interactions between CD40LG^+^ CD4^+^ T cells and CD40^*+*^ DCs in the CRT cohort (**Figure 2B**; **Methods**). These results are consistent with pre-clinical and clinical reports that CRT can induce immunogenic cell death, which increases tumor antigen availability, and stimulates the production of type I interferons, in turn activating DCs away from regulatory/suppressive cDC2, pDC and mregDC states towards improved capacity for cross-presentation (*17*–*22*),. Moreover, combinations of agonistic CD40 antibodies with chemotherapy and PD-1 inhibition has substantial benefit in some patient groups (*64*).

Intrinsic gene expression levels in immune cells differed as a function of treatment status (**Methods**), even in subsets whose proportions were comparable. For example, following CRT, CD8^+^ T lymphocytes expressed markers of altered differentiation (*e*.*g*., *SLAMF6, CD69, STAT4, IL7R*, shift from *ITGAE* to *ITGA4*) and TCR signaling (*e*.*g*., *ITK* and *FYN)* (**Figures 2D, S5A; Table S2**). While most well-established immune checkpoint receptors were not differentially expressed, *ENTPD1* (CD39) was marginally lower in the treated group (**Table S2**). In contrast, both inhibitory (*CD96*) and activating (*CD226*) members of the TIGIT/CD155 immune checkpoint family (*65*) were higher in the post-treatment context. In macrophages, there was a treatment-associated lower expression of genes for the uncommitted state M0 (*47, 66*) (*e*.*g*., *MARCO, AQP9, CYP27A1, NCF2*; **Figures 2D, S5B; Table S2**), and higher expression of macrophage polarization markers representing both the classical “pro-inflammatory” state M1 (*CD86, TNFSF8, TNFRSF1B, IFI44L, LILRB2*) and alternative “tissue repair” program M2 (*CD163, MRC1, TGFB1, TGFBI, TGFBR2, SIGLEC1, MS4A6A, FES*) (*47, 67*–*69*). CRT was also associated with higher expression of MHC class II (MHC-II) (*CD74, HLA-DPA1, HLA-DPB1*) and TGF-β pathway genes in macrophages (**Figures 2D, S5B**). Concomitantly, the lower expression of the TGF-β pathway in CAFs from treated tumors (**Figure S11A**) suggests a potential myeloid-specific immunosuppressive mechanism and may partly explain the efficacy of the TGF-β modulator, losartan, in combination with neoadjuvant CRT (*6, 70*).

### Interferon Signaling and Immune-Promoting Responses in Malignant Cells after Treatment

Analysis of genes differentially expressed by malignant cells in untreated *vs*. treated tumors (**Figure 2C**) revealed elevated expression of type I and II interferon (IFN) response genes following neoadjuvant CRT. This was detected even when cells from the two patients with germline *BRCA2* mutations were removed from the treated group (PDAC_T_1, PDAC_T_2; **Figure S5C; Table S1**). This phenomenon could be due to a release of damage associated molecular patterns (DAMPs) and pattern recognition receptors (PRRs) by CRT that converge on interferon signaling (*71*–*73*).

Notably, the IFN-γ pathway genes that are elevated in the malignant cells post treatment (**Figure 2C**) appear to be associated with immune-promoting rather than immune suppressive programs. None of the nonclassical MHC class I (MHC-I) genes that are thought to be immunosuppressive were differentially overexpressed post-treatment; the classical *HLA-B* was the only MHC-I molecule that was higher in the post-treatment context (**Figure 2C**) (*23, 74*). Furthermore, inferred interactions between *HLA-A, -B*, or *-C* on malignant cells and *CD3G* on CD8^+^ T-cells were higher post-treatment (**Figure 2B**). These data suggest that the addition of type I interferons may improve PDAC outcomes, although adjuvant CRT plus interferon alpha 2b did not improve survival compared to chemotherapy (*75, 76*). Overall, these results are consistent with reports that CRT may enhance MHC-I expression for recognition by cytotoxic immune cells (*77*).

### Subtype-Specific Treatment Response in Malignant Cells suggests that Basal-like Malignant Cells are more Resistant to CRT than Classical-like Cells

The differentially expressed genes in malignant cells suggest a relative increase in basal-like cells and a decrease in classical-like cells in treated *vs*. untreated tumors. Genes differentially overexpressed in malignant cells from post-treatment tumors were enriched in the basal-like signature (*9, 78*) (p = 2.05 × 10^−6^, hypergeometric test), and included *TP63*, a master transcription factor for the PDAC squamous subtype (*79*) (**Figure 2C**). Conversely, differentially underexpressed genes were enriched for the classical-like signature (p = 4.46 × 10^−24^), including the hallmark transcription factor *GATA6* (*7, 9, 78, 80*) (**Figure 2C**). Consistently, human PDAC cell lines become enriched for the basal-like subtype after FOLFIRINOX (*14, 81*). These observations could reflect either cell state changes following CRT, differential sensitivity to CRT, or both. Notably, the response to first-line combination chemotherapy for advanced PDAC is significantly better in patients with *GATA6*-expressing classical subtype tumors, supporting a model of increased sensitivity of classical-like cells to CRT.

Furthermore, genes differentially expressed *within* the treated cohort between malignant cells from patients with high (>10%) *vs*. low residual neoplastic content (**Figure S5D; Tables 1-2**) were also associated with the basal-like *vs*. classical-like distinction. Specifically, genes elevated in the high residual group were enriched for the basal-like signature (p = 8.34 × 10^−8^, hypergeometric test), and those underexpressed were enriched for the classical-like subtype (p = 1.65 × 10^−51^) (**Figure S5D**). Interestingly, *MUC16*, a source of neoantigens when mutated, and a potential target for chimeric antigen receptor (CAR) T cell therapy (*82*), was elevated in high-residual tumors. This may provide rationale for testing immunomodulatory neoadjuvant strategies to complement CRT. Overall, this analysis supports a model of a subtype-dependent response to CRT, such that basal-like neoplastic cells are more resistant to cytotoxic therapy than classical-like neoplastic cells. Future studies of matched pre- and post-treatment tumors will help distinguish direct selection on existing cell states from treatment-induced state changes.

In malignant cells, treatment was also correlated with distinct expression of genes needed to maintain the Wnt/β-catenin niche, which is crucial to treatment resistance, and can be mediated by either paracrine interactions with CAFs or autocrine signaling by malignant cells (*83*). The expression of the Wnt family receptor *LRP5* was sustained in surviving malignant cells post-treatment, but there appeared to be shifts in the source of Wnt signaling. CRT was associated with a concomitant differential underexpression in *WNT5A* by CAFs and overexpression in autocrine *WNT7B* signaling by malignant cells (**Figures 2B-C**), which has previously been shown to drive anchorage-independent growth and worse disease-specific survival in PDAC (*84*).

### Novel Malignant Cell Programs Reveal a Refined Molecular Classification

Next, we sought to better characterize expression states within the malignant cells across patients. Consistent with recent reports of intra-tumoral subtype heterogeneity, nearly every untreated tumor contained both basal-like and classical-like cells (**Figure S7; Methods**) (*78, 85*), with the two states being largely mutually exclusive. In the treated cohort, bulk signatures overlapped in the same nucleus, suggesting that the basal-like and classical-like signatures derived in the treatment-naïve setting may be less relevant in the neoadjuvant treatment context. This further supports the possibility of state changes and highlights the need to identify *de novo* molecular subtypes following CRT.

Despite substantial inter-tumor heterogeneity (**Figures 1C, S1**), we learned recurrent gene expression programs across malignant cells of different tumors by consensus non-negative matrix factorization (cNMF). We performed cNMF separately for the untreated and treated malignant cells and focused on the programs shared between patients that were biologically distinct (**Figures 3A-B; Tables S3-4; Methods**) (*86*). We annotated each program based on its top 200 weighted genes (**Methods**). In both untreated and treated tumors, we identified nine malignant programs that reflected either their lineage or cell state, though there was partial overlap among them (**Figures 3A-B**).

**Figure 3.**
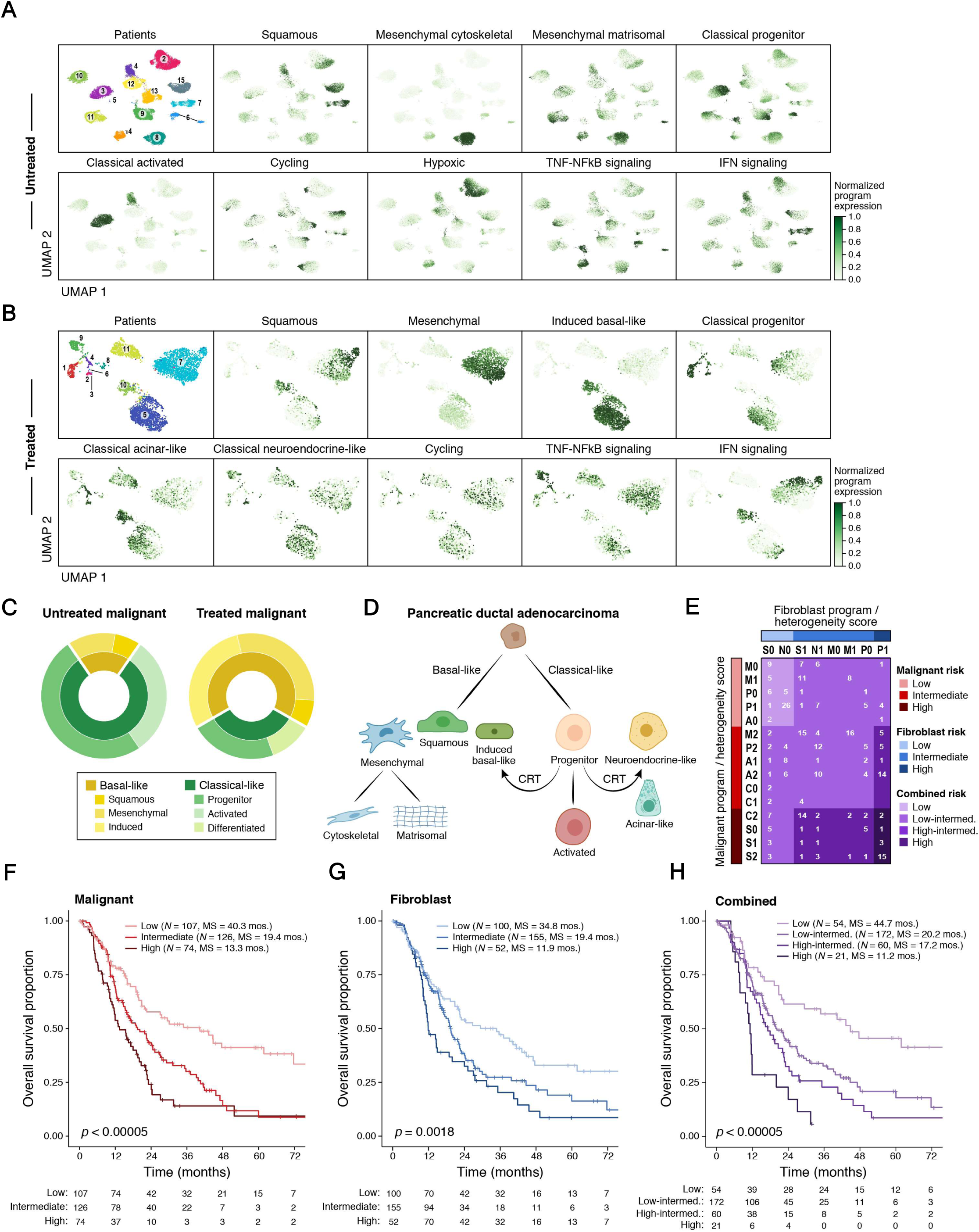
Refined Molecular Taxonomy of PDAC Improves Clinical Prognostication and Highlights a Shift from Classical-like to Basal-like and Differentiated-like Programs in Malignant Cells. (A,B) A consensus NMF (cNMF) expression program dictionary in untreated and treated tumors. UMAPs of single nucleus profiles (dots) from untreated (A) and treated (B) tumors, colored by patient (top left) or by the score derived for each cell-cNMF program pair (**Methods**). (C) Shift towards basal-like programs in CRT treated tumors. Proportion of malignant cells primarily expressing each lineage-specific malignant cell cNMF program within untreated (left) and treated (right) tumors. (D) Refined PDAC molecular taxonomy with proposed model of transcriptional programs and their relationships. (E-G) Refined molecular taxonomy of malignant and fibroblast cells has prognostic value. (E) Tumor assignment to risk categories defined by primary snRNA-seq program (first letter) and heterogeneity score (second number) for malignant cells (rows) and fibroblasts (columns) separately (red and blue shades, **Methods**) and by their combination (purple shades, **Methods**) for bulk RNA-seq classification of patients with untreated resectable PDAC (n = 307). The number of tumors in each stratification is listed. For malignant programs: P = *classical progenitor*, A = *classical activated*, M = *mesenchymal matrisomal*, C = *mesenchymal cytoskeletal*, S = *squamous*. For fibroblast programs: N = *neurotropic*, S = *secretory*, P = *mesodermal progenitor*, M = *myofibroblast*. For heterogeneity score: 0 = fewer than two highly-expressed programs, 1 = two highly-expressed programs (malignant) or two or more high-expressed programs (fibroblast), and 2 = three or more highly-expressed programs (malignant). (F-H) Kaplan-Meier survival analysis of bulk RNA-seq cohort (n = 307) based on risk groups from malignant (F), fibroblast (G) and combined (H) strata, as defined in (E). Survival distributions are compared by the log-rank test. Number of patients at risk at the beginning of each time interval is shown in the table.

In untreated tumors, there were five lineage-specific programs: three spanned basal-like phenotypes involving the epithelial-mesenchymal transition (*87, 88*) (*squamous, mesenchymal cytoskeletal, mesenchymal matrisomal*), with the *squamous* program closely overlapping the basal-like A subtype (*78*); two spanned classical-like phenotypes (*classical progenitor, classical activated*) (**Figure 3A**) with *the classical progenitor* program containing transcription factors (TFs) involved in endoderm lineage development (*HNF1A, DLX2, PRDM6*, and *FOXO4*) and the *classical activated* program also containing genes involved in secretion, cell polarization, and cytoskeletal remodeling. The remaining four cell state programs were *cycling, hypoxic, TNF-NFkB signaling*, and *interferon signaling*. In treated tumors, there were six lineage-specific programs (*squamous, mesenchymal, induced basal-like, classical progenitor, classical acinar-like, classical neuroendocrine-like*) and three cell state programs (*cycling, TNF-NFkB signaling, interferon signaling*) (**Figure 3B**).

Programs varied in the extent to which they co-occurred within the same cell and associated with one another. In both untreated and treated tumors, *IFN signaling* and *squamous* program scores were correlated across nuclei (**Figure S8**), as were those of *mesenchymal* with *TNF-NFkB signaling. TNF-NFkB signaling* was also correlated with the *induced basal-like* and *classical neuroendocrine-like* programs in the post-treatment context. Thus, basal-like cells may be more inflammatory/immunogenic overall than classical-like cells. Taken together, the increase in expression of basal-like genes, interferon signaling, and immune response promoting genes (**Figure 2C**) in treated *vs*. untreated samples may reflect coupled programs in the same individual cells (**Figure S8**), which could in turn drive immune cell state changes (**Figures S5A-B**). One possibility is that these are driven through the effects of p63, especially the ΔNp63 isoform, though this remains an area of controversy in the field (*11, 89, 90*).

### A Shift from Classical-like to Induced Basal-like or Terminally-Differentiated Pancreatic Cell Programs may Contribute to Resistance Following Treatment

Compared to the untreated group, post-treatment malignant cells scoring highly for basal-like programs were enriched, while those scoring highly for classical-like programs were depleted (66% vs 19%; p = 0.0001, Fisher’s exact test; **Figures 3C, S7**). This was consistent with our differential expression analysis (**Figure 2C**). Despite the overall reduction of classical-like cells post-treatment, the classical-like programs present in residual cells spanned a broader range of pancreatic cell lineages: *classical progenitor* (*GATA6* TF), *classical acinar-like* (endoderm lineage

TFs and characteristic digestive enzymes), and *classical neuroendocrine-like* (enriched for neural genes and target genes of pancreatic neuroendocrine TFs such as *HNF3* and *NKX6*). This suggests that neoadjuvant CRT either selects for or drives cells towards a state of increased pancreatic differentiation (**Figure 3B**) and mirrors the relative enrichment in non-neoplastic acinar and endocrine cells after CRT (**Figures 2A, S4**).

In addition, the post-treatment *induced basal-like* program shared features of both classical-like (*classical activated*) and basal-like (*squamous*) programs, along with *MUC16* (also a member of the untreated *interferon signaling* program, **Figure 3A**) and genes associated with chemoresistance (*MUC13, CEACAM6, MUC5, FGF19, ABCC3, TNFAIP2, KLK11, DUSP1, OLFM4, AQP5*)(*91*– *101*) (**Figure 3B**). This *induced basal-like* post-treatment program may reflect a phenotypic shift from the more therapeutically vulnerable classical-like subtype to the more resistant basal-like state reminiscent of recent *ex vivo* observations (*81*).

More generally, these programs may reflect a shift towards either basal-like or terminally-differentiated pancreatic cell states that are advantageous for surviving CRT compared to a less differentiated classical-like phenotype (**Figure 3D**). This model is consistent with *in vitro* and *in vivo* evidence for lineage plasticity in normal pancreatic ductal progenitor cells (*102*) and differentiated pancreatic cells—including a role for MYT1/MYT1L, a member of the *classical neuroendocrine-like program*, in ductal-neuroendocrine reprogramming (**Figure 3B; Tables S3-4**) (*103*–*108*).

### Treatment-Associated Contractile Phenotype in Cancer-Associated Fibroblasts

Because fibroblasts have emerged as a key instigator of tumor-immune evasion and therapeutic target (*109*), we leveraged the successful capture of fibroblasts in our data to assess their subsets, programs, and response to treatment. Although three signatures were recently reported in a scRNA-seq study of human and murine PDAC (myofibroblastic CAFs (myCAF), inflammatory CAFs (iCAF), and antigen-presenting CAFs (apCAF)) (*51*), these did not adequately segregate our snRNA-seq data, likely reflecting the substantial underrepresentation of CAFs in earlier scRNA-seq studies (*51*) (**Figure S10**). We thus separately applied cNMF to learn four snRNA-seq programs each for the untreated and treated CAFs (**Figure S11B**; **Table S5**; **Methods**).

Three of the programs were shared between untreated and treated CAFs: *myofibroblast, neurotropic*, and *secretory*, while a *mesodermal progenitor* program (mesodermal developmental genes and TFs) was discovered only in untreated CAFs, and a *neuromuscular* program (muscle development, contractility, synapse, and action potential genes) was only learned in treated CAFs (**Figure S11B**). The *myofibroblast* program included extracellular matrix, motility, and wound response genes; the *neurotropic* program featured neural development, synaptic, and axonal guidance genes; and the *secretory* program encompassed protein targeting, secretory vesicle, exocytosis, and cytokine signaling genes including modulators of diverse immune functions (e.g., *CXCL14, LGALS1, CST3, PPIA, LGALS3BP, CD59, CD81*, and *IFITM3*) (*110*–*115*). In untreated CAFs, the *myofibroblast* and *mesodermal progenitor* programs were positively correlated (**Figure S11C**).

The marked enrichment of the *myofibroblast* phenotype after neoadjuvant CRT (**Figure S11D**) may be consistent with an induction of genes associated with muscle development and contraction after CRT seen in differential expression analysis (**Figure S11A**). *BCL2*, an anti-apoptotic protein, was also expressed at a higher level in post-CRT myofibroblasts (**Figure S11A**). *BCL2* inhibition with Navitoclax (ABT-263) induced apoptosis of myofibroblasts and impeded tumor growth in a xenograft mouse model of hepatocellular cancer (*116*), and could thus be a relevant therapeutic strategy in the neoadjuvant setting. Conversely, the expression of *FAP*, a member of the *mesodermal progenitor* program, was significantly reduced after CRT, suggesting that depletion of CAFs by blocking FAP may not confer added value, despite prior studies showing promising anti-tumor effects in some model systems (*27, 117*) (**Figure S11A**). Treatment was also associated with higher *CXCL12* expression in CAFs (**Figure S11A**) and increased predicted *CXCL12*-*CXCR4* interactions between CAFs and CD8^+^ T-cells (**Figure 2B**), which has been linked to inhibition of T-cell migration to the TME (*118*). Moreover, CXCR4 inhibition improves PDAC sensitivity to anti-PD-1/PD-L1 immunotherapy (*119, 120*). Thus, modulating this axis may further improve clinical outcomes associated with neoadjuvant CRT.

**Clinically Relevant Molecular Taxonomy Based on Malignant and Fibroblast Programs Improves Prognostication**

Prior survival analyses that stratified patients by bulk expression subtypes only discerned binary prognostic differences between basal-like <u>and</u> classical-like tumors (*7, 10*), but finer subsets, such as three non-basal bulk subtypes (*10*) (pancreatic progenitor, immunogenic, ADEX) were indistinguishable. We used our *de novo* snRNA-seq programs for untreated malignant cells and fibroblasts to stratify bulk RNA-seq profiles from untreated, resected primary PDAC in the TCGA (*11*) and PanCuRx (*78*) cohorts (n = 307; **Methods**). To account for the effects of different cell types on survival, we considered both malignant and fibroblast untreated programs, separately and in combination, to assign patients to risk categories (**Methods**). Briefly, we scored each untreated tumor by the five lineage malignant programs and four fibroblast programs. Each tumor was characterized by (1) its top scoring program (primary program) and (2) the number of highly expressed programs (heterogeneity score) and assigned by these two criteria to one of 15 possible malignant classes (**Figure 3E**, rows) and 8 possible fibroblast classes (**Figure 3E**; columns). Next, we inspected the survival curves associated with each class (**Figure S9**) and aggregated them into three putative risk categories (separately for malignant or fibroblast): low, intermediate, and high (**Figure 3E**, red and blue color bars). Finally, we also assigned tumors to four combined risk strata by integrating the malignant and fibroblast risk groups: low (both low), high (both high), high-intermediate (one high, one intermediate), low-intermediate (all others) (**Figure 3E**, internal grid, purple color code).

Kaplan-Meier (KM) analyses of overall survival (OS) stratified by either primary program, heterogeneity score, or both were separately prognostic for malignant cells and fibroblasts (**Figures S9A-F**). Among the malignant programs, *mesenchymal matrisomal* and *classical progenitor* were associated with the best OS, *classical activated* was associated with intermediate OS, and *squamous* and *mesenchymal cytoskeletal* were associated with the worst OS (**Figure 9A**). Notably, the basal-like *mesenchymal matrisomal* program had survival outcomes comparable to *classical progenitor*. Moreover, we identified the *classical activated* program as a classical-like subset with worse outcomes than the *classical progenitor* program. Among fibroblast programs, the *secretory* and *neurotropic* programs were associated with longer survival while the *myofibroblast* and *mesodermal progenitor* programs were associated with shorter survival (**Figure S9D**). These findings were consistent with prior work associating myofibroblasts and mesenchymal stem-like cells with poor prognosis in a range of solid cancers (*121*–*123*). For both malignant cells and CAFs, an increasing number of highly scoring programs in one tumor (heterogeneity score) was associated with worse OS (**Figures S9B**,**E**), consistent with the known association between intratumoral heterogeneity, treatment resistance, and poor outcomes (*29, 124*). When combining the malignant and fibroblast risk groups into a four-tier risk stratification (**Figure 3E**), there was a significant prognostic difference among the combined strata (log-rank p < 0.00005; **Figure 3H**) with a greater dynamic range (median survival: 11.2 to 44.7 months) than seen with prior classifications (*10*).

### Digital Spatial Profiling Reveals Malignant- and Fibroblast-Intrinsic Programs May Modulate Local Immune Microniches

Our snRNA-seq analysis highlighted multiple potential inter-compartmental interactions among malignant, stromal and immune cells, including those associated with CRT. Identifying cancer cell- and fibroblast-intrinsic programs that may govern local immune microniches remains an open question in PDAC research, with prior studies disagreeing on whether the basal-like or classical-like subtype is correlated with immune exclusion (*11, 89, 90*). Moreover, elucidating the relationships between our newly-identified neoplastic, lineage-specific programs (*81, 102*–*108*) and local immune infiltration will be critical in guiding therapeutic development.

To address this challenge, we performed digital spatial profiling (DSP) with the GeoMx platform (NanoString) and a cancer transcriptome atlas (CTA) probe set. In this method, UV-photocleavable barcode-conjugated RNA ISH probes against 1,412 target mRNAs were used to capture and profile mRNA from user-defined regions of interest (ROI) (**Figure S12A; Methods**) (*125*). A four-color immunofluorescence slide scan for each specimen (**Figure 4A**) showed intra-tumoral diversity in tissue architecture, allowing us to profile three distinct classes of ROIs encompassing neoplastic cells with either (1) immune and CAF infiltration, (2) immune infiltration only, or (3) CAF infiltration only (**Figure 4B**). We then used custom illumination masks to separately capture RNA from areas of illumination (AOI) enriched for one cell type within the ROI, collected RNA ISH barcodes from each AOI in a spatially-indexed manner, and counted transcripts by sequencing (**Figure S12A; Methods**).

**Figure 4.**
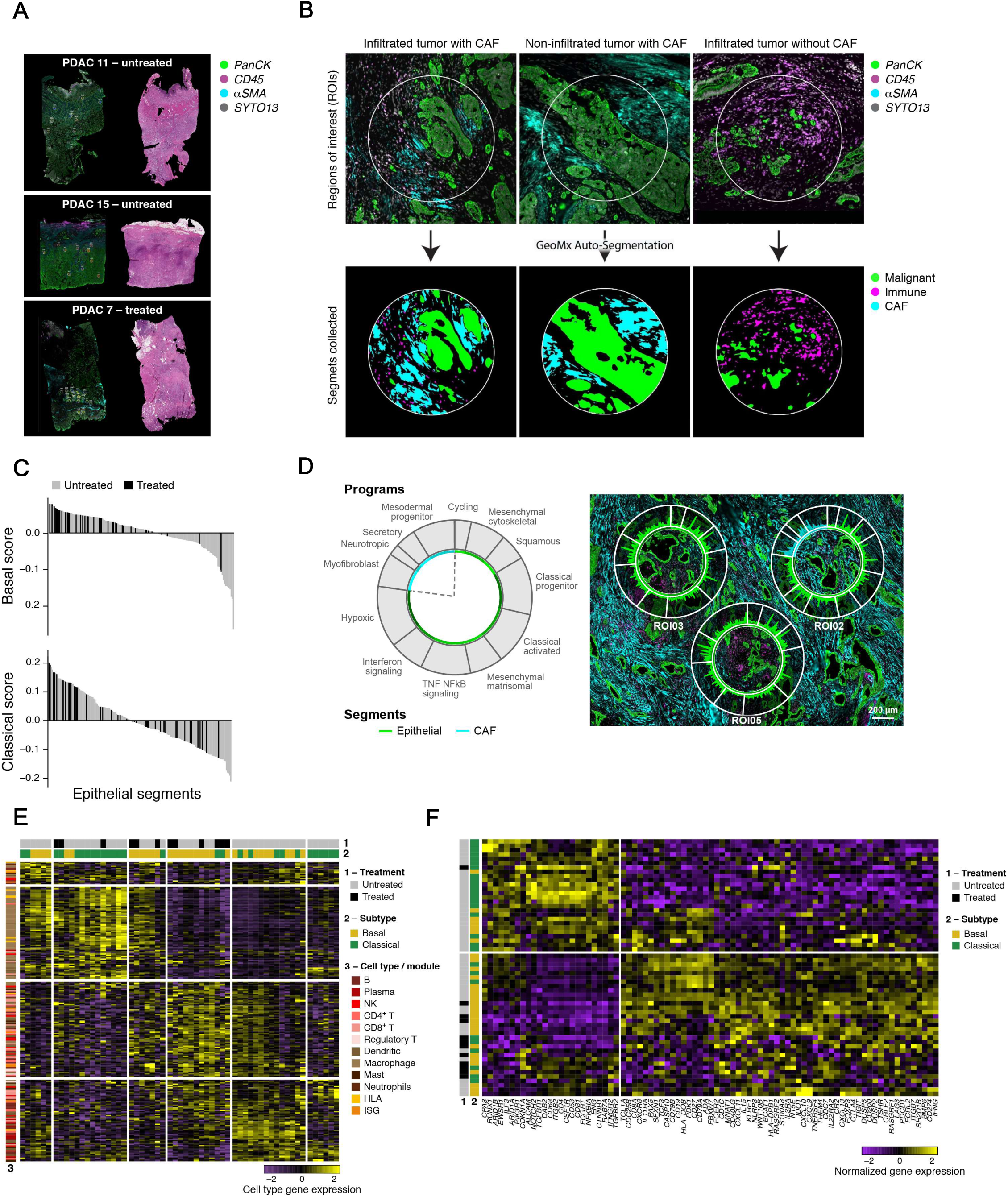
Basal-like and Classical-like Programs are Associated with Spatial Niches with Distinct Quantity and Quality of Immune Infiltration. (A,B) Definition of distinct regions of interest (ROIs) with GeoMx DSP. (A) Immunofluorescence images (left, GeoMx DSP) and matched consecutive hematoxylin and eosin (H&E)-stained FFPE sections (5 µm thickness, right) of three patients (labeled, top), showing the selected ROIs (circles). (B) Top: GeoMx DSP immunofluorescence images with selected ROIs (circles, 600 µm diameter) representing three classes of epithelial niches infiltrated with either both immune cells and CAFs (left), only with CAFs (middle), or only with immune cells (right). Bottom: Segmentation masks on the ROIs, used to enrich for the epithelial, CAF, and immune compartments. Gray = SYTO13 (nuclear stain), green = anti-panCK, magenta = anti-CD45, cyan = anti-αSMA. (C) Increased basal-like programs in treated tumors *in situ*. Aggregate basal-like and classical-like signature scores (*y* axis) for each epithelial area of interest (AOI) from untreated (grey) and treated (black) tumors, rank-ordered by score as determined from the GeoMx Cancer Transcriptome Atlas (CTA). (D) GeoMx Whole Transcriptome Assay (WTA) also detected intra-tumoral heterogeneity in untreated malignant and CAF programs. Magnitude of expression (amplitude) for each gene (position in circle plot) in each malignant or CAF program (segment in circle plot, schematic on left) in three ROIs selected and segmented as for the CTA in (B) on specimen PDAC_U_7. (E) Basal-like and classical-like malignant programs associate with immune niches with distinct characteristics. Expression (z-score of normalized counts across AOIs; color bar) of immune cell signature genes (rows) from diverse cell types and signatures (E, color legend and left bar) across immune AOIs (columns) from untreated (grey) and treated (black) tumors in ROIs with either basal-like (yellow) or classical-like (green) malignant cells. HLA = human leukocyte antigen module, ISG = interferon-stimulated gene module. (F) Expression (z-score of the normalized counts across AOIs; color bar) of subtype-associated immune cell type genes (rows) across immune AOIs (columns) from untreated (grey) and treated (black) tumors in ROIs with either basal-like (yellow) or classical-like (green) malignant cells.

We used our snRNA-seq cell type signatures to deconvolve the spatial profiles. The malignant, CAF, and immune AOIs clustered appropriately by cell type, demonstrating the coherence and complementarity of the two platforms (**Figure S12B**). We also mapped the expression of each malignant and CAF program onto the spatial data (**Figures 3A-B, S11B**). Consistent with the snRNA-seq results, within the malignant compartment, CRT was associated with a higher basal-like score (mixed effect model p = 0.0035) but not classical-like score (mixed effect model p > 0.2) compared to untreated specimens (**Figure 4C**). Because only 269 genes were shared between the 1,412 gene CTA panel and the 2,037 program genes, we also profiled three untreated specimens using a whole transcriptome atlas (WTA; 18,269 genes; one probe per gene; 1,918 of the program genes) (**Figure 4D**; **Methods**). Programs from untreated patients had concordant expression within the WTA and CTA data (Spearman’s ρ = 0.54-0.999; **Figure S12E**) and mapped to locally-defined architecture within the tumor in a comparable manner (**Figure 4D**).

Relating the expression of each malignant and CAF program (**Figures 3A-B, S11B**) to the physical architecture of the tumor (**Methods**), the CAF *myofibroblast* program was associated with immune exclusion in treated samples (mixed effect model p = 0.021), consistent with the role of myofibroblasts in mediating an immunosuppressive TME (*126*) (**Figure S11B**). Immune-excluded ROIs (class 3) were also associated with higher untreated *classical activated* malignant program expression (mixed effect model p = 0.020), while the untreated *mesenchymal matrisomal* malignant program was associated with immune infiltration (mixed effect model p = 0.0035) (**Table S6**). The association between the *classical activated* program and immune exclusion may be related to the poorer prognosis we found for this program in untreated patients compared to other classical-like programs (**Figure 3E-F**). Similarly, the association between the *mesenchymal matrisomal* program and immune infiltration may be related to the longer survival we found for this program in untreated patients compared to other basal-like programs (**Figure 3E-F**). In post-treatment tumors, the *classical neuroendocrine-like, squamous*, and *induced basal-like* malignant programs associated with or trended towards an association with immune-infiltrated ROIs (mixed effect p = 0.029, 0.074, 0.091, respectively). These spatially resolved findings support an overall association between basal-like programs and immune infiltration and conversely, between specific classical-like programs and immune exclusion with certain exceptions. Despite these overall trends, some ROIs that contained high classical scoring epithelial segments also featured an immune infiltrate, which warranted further exploration.

There were also differences in the types of immune infiltrates surrounding basal-like or classical-like malignant cells. We observed the association between immune infiltrates with malignant cell subtype by unsupervised clustering of immune cell type-specific and functional module genes measured within immune AOIs (p = 0.0339, χ^2^ test; **Figure 4E**; **Methods**). Moreover, immune AOIs in classical-like segments had higher expression of clusters of macrophage-rich genes (p = 0.0341, t-test), while those from basal-like segments had higher expression of clusters of T lymphocyte, B lymphocyte and dendritic cell genes (and depleted of macrophage genes) (p = 6.74 × 10^−5^ and 0.047, t-test, for macrophage-depleted gene clusters A and B, respectively, **Figure 4E**; **Figure S12C**). Moreover, at the individual gene level, epithelial AOI subtype (classical-like vs basal-like) was associated with expression of distinct immune lineage-restricted and modulatory genes (**Figure 4F**; **Table S7)**: basal-like segments had immune AOIs with higher expression of *IFNG-*induced chemokines (*CXCL9, CXCL10, CXCL11*), markers of cytotoxic T lymphocytes and NK cells (*CD8A, CD3E, CD247, GZMA, GZMB, GZMK, PRF1, NKG7, NCAM1*), immune checkpoints/markers of cytotoxic T lymphocyte exhaustion (*PDCD1, CD274, LAG3, ENTPD1, TIGIT, IDO1*), and markers of regulatory T cells (*NT5E, FOXP3*) (**Figure 4F**). In contrast, classical-like segments had immune AOIs with higher expression of MHC-II (*HLA-DQA1/2, HLA-DOA, HLA-DMA, HLA-DPA1*) and macrophage markers (*CD68, CD163, CSF1R*). Taken together, these analyses suggest distinct immune niches associated with basal-like and classical-like malignant cells.

## DISCUSSION

Leveraging single-nucleus RNA-seq of frozen archival PDAC, we comprehensively identified common biological programs among untreated and post-CRT malignant cells and cancer-associated fibroblasts. This refined molecular taxonomy of PDAC (**Figure 3D**) allowed us to stratify patients from bulk RNA-seq profiles of their tumors into prognostic risk groups defined by malignant cell and fibroblast program scores. We detected differences in malignant and stromal cell composition and programs following treatment, including an enrichment of basal-like and differentiated states in malignant cells and contractile phenotypes in CAFs. Integrating the snRNA-seq and spatial profiling data from the same tumors, we demonstrated how these differences associate with distinct immune microniches.

Although our study does not include matched pre- and post-treatment specimens and the treated cohort size is modest and cannot be stratified by treatment regimen, our data refine and clarify the overarching distinction between basal-like and classical-like programs and how they are affected by CRT. We identified squamous and mesenchymal subclasses within the basal-like subtype in both the untreated and treated contexts, as well as bi-lineage differentiated classical-pancreatic states and an induced basal-like phenotype that arise in the treated setting. Collectively, our analysis suggests that CRT may drive a shift towards basal-like and differentiated classical phenotypes and away from classical progenitor states, which may be due to a combination of direct selection on pre-existing states and induced plasticity (**Figures 2C**,**3D**). The basal-like programs are also intimately connected to immune-activating interferon and TNF-NFkB programs in the malignant cells (**Figure S8**), and their enrichment in the CRT context may be conducive to a more activated, immunogenic environment. Thus, at least some of the CRT-resistant cell states enriched or induced by CRT may yield a tumor microenvironment that is more susceptible to some immunotherapies.

Our spatially-resolved transcriptomics analysis further supported the hypothesis that basal-like malignant cell programs may facilitate a greater degree of immune infiltration compared with classical-like programs (*89*). Moreover, the immune infiltrates associated with basal-like and classical-like malignant cells were distinct, suggesting potential strategies for differentially targeting these phenotypes, with immune checkpoint inhibitors for the former and myeloid-directed therapies such as CD40 agonists and TGF-beta modulators (*e*.*g*., losartan) for the latter. Similar phenomena have been observed in other cancer types such as breast, in which the triple-negative subtype has enhanced basal-like features compared to others and is similarly associated with elevated immune cell infiltration that correlates with greater response to immune checkpoint inhibitors (*127, 128*).

Our snRNA-seq data also helps address the open question of whether the previously identified exocrine-like and ADEX subtypes of PDAC truly exist or if they represent normal tissue contamination (*7, 8, 10, 11, 129*–*131*). We confirmed the existence of both exocrine- and endocrine-like cancer cells (with inferred CNAs) in post-treatment but not untreated tumors. Thus, it is plausible that endoderm-differentiated cancer cell phenotypes are only prevalent enough to be detected under treatment selection pressure, and may alternatively reflect normal cell contamination in treatment-naïve bulk studies.

The presence of relatively resistant neuroendocrine- and exocrine-like cancer cells after neoadjuvant CRT is also clinically important. Neuroendocrine cells in PDAC and precursor lesions have been shown to promote tumorigenesis via neuronal cross-talk, and may thus be partly responsible for the enrichment in Schwann cells associated with treatment (**Figure 2A**) (*130, 132*). Moreover, primary PDAC cell lines from xenografts featuring the *HNF1A*-expressing exocrine subtype (*130*) are relatively resistant to small molecule tyrosine kinase inhibitors (TKIs), in a manner dependent on an inducible *CYP3A5* that oxidizes and inactivates the TKIs. Notably, *CYP3A5* is a member of some of the classical-like topics in our cohort, but *CYP3A5* and its regulator *HNF4A* were repressed in both the post-CRT cohort and its high-residual post-CRT subgroup (**Figures 2C, S5D**; **Table S2**). *NF1I2*, which modulates drug-induced *CYP3A5* upregulation is also repressed in the high-residual post-CRT subset. Taken together, these results suggest that CRT may sensitize residual basal-like cancer cells to TKIs and warrants further investigation.

Overall, our study provides a high-resolution molecular framework for understanding the intra-tumoral diversity of pancreatic cancer and treatment-associated changes, spatial associations among malignant/fibroblast-intrinsic programs and both quantitative and qualitative differences in immune microniches, and clinically-relevant prognostication. These findings can be harnessed to augment precision oncology efforts in pancreatic cancer.

## METHODS

### Human Patient Specimens

For inclusion in this study, patients had non-metastatic pancreatic ductal adenocarcinoma and went to surgical resection with or without neoadjuvant radiotherapy and/or chemotherapy. Some patients received additional neoadjuvant therapy in the form of immune checkpoint inhibitors or losartan, an angiotensin II receptor type 1 antagonist. All patients were consented to protocol 2003P001289 (principal investigator: CFC; co-investigators: ASL, WLH), which was reviewed and approved by the Massachusetts General Hospital (MGH) Institutional Review Board. Resected primary tumor samples were examined to confirm neoplastic content by a board-certified pathologist (MMK) and then snap frozen and stored at −80°C for up to 7 years prior to processing. Specimens were screened for an RNA integrity number (RIN; Agilent RNA 6000 Pico Kit, cat. No. 5067-1513) greater than an empirically determined threshold of 6; only specimens with RIN > 6 were processed further.

**Nucleus Isolation from Frozen Samples**

We have recently published a toolbox for snRNA-seq of tumors spanning a broad range of nucleus isolation techniques for various tissue/tumor types (*43*), but not PDAC. The following protocol is an adaptation and optimization of this prior work specifically for the unique tissue requirements of pancreatic tumors. A 2x stock of STc buffer in nuclease-free water was prepared with a final concentration of 292 mM NaCl (ThermoFisher Scientific, cat. no. AM9759), 40 mM Tricine (VWR, cat. no. E170-100G), 2 mM CaCl_2_ (VWR, cat. no. 97062-820), and 42 mM MgCl_2_ (Sigma Aldrich, cat. no. M1028). For each specimen, 2 mL of NSTcPA buffer was prepared by combining 1 mL of 2x STc buffer, 40 µL of 10% Nonidet P40 Substitute (Fisher Scientific, cat. no. AAJ19628AP), 10 µL of 2% bovine serum albumin (New England Biolabs, cat. no. B9000S), 0.3 µL of 1M spermine (Sigma-Aldrich, cat. no. S3256-1G), 1 µL of 1M spermidine (Sigma-Aldrich, cat. no. S2626-1G), and 948.7 µL of nuclease-free water. For each specimen, 3 mL of 1x working STc buffer was made by diluting 2x STc 1:1 in nuclease-free water.

NSTcPA buffer (1 mL) was pipetted into one well of a 6-well plate (Stem Cell Technologies, cat. no. 38015) on ice. The frozen tumor specimen was removed from −80°C and placed in a petri dish on dry ice. Using a clean razor blade, the desired regions of the tissue were cut while the specimen remained frozen (at least 10-20 mg). The remainder of the specimen was returned to −80°C for subsequent use. The selected tissue was transferred into the NSTcPA buffer and manually minced with fine straight tungsten carbide scissors (Fine Science Tools, cat. no. 14568-12) for 8 minutes. The homogenized tissue solution was then filtered through a 40 µm Falcon cell filter (Thermo Fisher Scientific, cat. no. 08-771-1) into a 50 mL conical tube. An additional 1 mL of NSTcPA buffer was used to rinse the well and filter. The total volume was brought up to 5 mL with 3 mL of 1x STc buffer and transferred into a 15 mL conical tube. The sample was spun for 5 min at 500xg, 4°C and the supernatant was removed. The pellet was resuspended in 100-200 µL 1x STc and then filtered through a 35 µm Falcon cell strainer (Corning, cat. no. 352235). Nuclei were quantified using a C-chip disposable hemocytometer (VWR, cat. no. 82030-468) and diluted in 1x STc as necessary to achieve a final concentration of 300-2,000 nuclei/µL.

### Single-Nucleus RNA-seq (snRNA-seq)

Approximately 8,000 nuclei per sample were loaded into each channel of a Chromium single-cell 3’ chip (V2 or V3, 10x Genomics) according to the manufacturer’s instructions. Single nuclei were partitioned into droplets with gel beads in the Chromium Controller to form emulsions, after which nucleus lysis, barcoded reverse transcription of mRNA, cDNA amplification, enzymatic fragmentation, and 5’ adaptor and sample index attachment were performed according to manufacturer’s instructions. Up to four sample libraries were sequenced on the HiSeq X Version 2.5 (Illumina) with the following paired end read configuration: read 1, 26-28 nt; read 2, 96-98 nt; index read, 8 nt.

### snRNA-seq Data Pre-Processing

BCL files were converted to FASTQ using bcl2fastq2-v2.20. CellRanger v3.0.2 was used to demultiplex the FASTQ reads, align them to the hg38 human transcriptome (pre-mRNA) and extract the UMI and nuclei barcodes. The output of this pipeline is a digital gene expression (DGE) matrix for each sample, which has quantified for each nucleus barcode the number of UMIs that aligned to each gene.

We filtered low-quality nuclei profiles by baseline quality control measures. First, we discarded profiles with fewer than 400 genes expressed or with greater than 20% of reads originating from mitochondrial genes. Additionally, we performed doublet detection over all nuclei profiles by using Scrublet (*133*) and removed all profiles with a Scrublet score greater than 0.2. To account for differences in sequencing depth across nuclei, UMI counts were normalized by the total number of UMIs per nucleus and converted to transcripts-per-10,000 (TP10K) as the final expression unit.

### Dimensionality Reduction, Clustering and Annotation

Following these quality control steps, treatment-naïve and neoadjuvant-treated specimens were aggregated into two separate datasets. The log_2_(TP10K+1) expression matrix for each dataset was used for the following downstream analyses. For each dataset, we identified the top 2,000 highly variable genes across the entire dataset using the Scanpy (*134*) *highly_variable_genes* function with the sample id as input for the batch. We then performed a Principal Component Analysis (PCA) over the top 2,000 highly variable genes and identified the top 40 principle components (PCs) beyond which negligible additional variance was explained in the data (the analysis was performed with 30, 40, and 50 PCs and robust to this choice). Subsequently, we built a *k*-nearest neighbors graph of nuclei profiles (*k* = 10) based on the top 40 PCs and performed community detection on this neighborhood graph using the Leiden graph clustering method (*135*). Distinct cell populations were identified and annotated using known cell type-specific gene expression signatures (*40, 47*–*49*). Individual nuclei profiles were visualized using the uniform manifold approximation and projection (UMAP) (*44*).

### Inferring Copy Number Aberrations from Single-Nucleus Profiles

InferCNV v3.9 (*136*) was run on all nuclei profiles for each tumor separately with a common set of high confidence non neoplastic cells used as the reference. We used a 100 gene window in sub-clustering mode and an HMM to predict the copy number aberration (CNA) count in each nucleus.

### Multiplexed Ion Beam Imaging (MIBI)

Formalin-fixed paraffin-embedded pancreatic tissue sections were cut onto gold MIBI slides (IONpath, cat. no. 567001) and stained at IONpath (Menlo Park, CA) with the internal Epithelial i-Onc isotopically-labelled antibody panel (IONpath): dsDNA_89 [3519 DNA] (1:100), β-tubulin_166 [D3U1W] (3:200), CD163_142 [EPR14643-36] (3:1600), CD4_143 [EPR6855] (1:100), CD11c_144 [EP1347Y] (1:100), LAG3_147 [17B4] (1:250), PD-1_148 [D4W2J] (1:100), PD-L1_149 [E1L3N] (1:100), Granzyme B_150 [D6E9W] (1:400), CD56_151 [MRQ-42] (1:1000), CD31_152 [EP3095] (1:1000), Ki-67_153 [D2H10] (1:250), CD11b_155 [D6×1N] (1:500), CD68_156 [D4B9C] (1:100), CD8_158 [C8/144B] (1:100), CD3_159 [D7A6E] (1:100), CD45RO_161 [UCHL1] (1:100), Vimentin_163 [D21H3] (1:100), Keratin_165 [AE1/AE3] (1:100), CD20_167 [L26] (1:400), Podoplanin_170 [D2-40] (1:100), IDO1_171 [EPR20374] (1:100), HLA-DR_172 [EPR3692] (1:100), DC-SIGN_173 [DCN46] (1:250), CD45_175 [2B11 & PD7/26] (3:200), HLA class 1 A, B, and C_176 [EMR8-5] (1:100), Na/K-ATPase_176 [D4Y7E] (1:100).

Quantitative imaging was performed using a beta unit MIBIscope (IONpath) equipped with a duoplasmatron ion source. This instrument sputters samples with O_2_^+^ primary ions line-by-line, while detecting secondary ions with a time-of-flight mass spectrometer tuned to 1-200 m/z+ and mass resolution of 1000 m/Δm, operating at a 100 KHz repetition rate. The primary ion beam was aligned daily to minimize imaging astigmatism and ensure consistent secondary ion detection levels using a built-in molybdenum calibration sample. In addition to the secondary ion detector, the MIBIscope is equipped with a secondary electron detector which enables sample identification and navigation prior to imaging.

For data collection, three fields of view were acquired for each sample by matching the secondary electron morphological signal to annotated locations on sequential H&E stained slides. The experimental parameters used in acquiring all imaging runs were as follows: pixel dwell time (12 ms), image size (500 μm^2^ at 1024 × 1024 pixels), primary ion current (5 nA O_2_^+^), aperture (300 μm), stage bias (+67 V).

### MIBI Image Processing, Segmentation and Quantification

Mass spectrometer run files were converted to multichannel tiff images using MIB.io software (IONpath). Mass channels were filtered individually to remove gold-ion background and spatially uncorrelated noise. HLA Class 1 and Na/K-ATPase signals were combined into a single membrane marker. These image files (tiff) were used as a starting point for single cell segmentation, quantification and interactive analysis using histoCAT (v1.76) (*137*). We followed a similar approach for segmentation as proposed for Imaging Mass Cytometry data (*137*–*139*). Briefly, we used Ilastik (*140*) to manually train three classes (nuclei, cytoplasm and background) to improve subsequent watershed segmentation using CellProfiler (*141*). Finally, the tiff images and masks were combined for histoCAT loading with a script optimized for MIBI image processing. All code, classifiers and configuration files are available at https://github.com/DenisSch/MIBI

### Differential Gene Expression Analysis

For each annotated cell type detected in both untreated and treated tumors, a differential gene analysis was performed between cells in the two populations to identify upregulated and downregulated genes. A Wilcoxon statistical test was used to compute the p-values for each gene and Bonferroni correction was applied to correct for multiple testing.

### Scoring Gene Signatures for Each Nucleus Profile

A signature score for each nucleus profile was computed as the mean log_2_(TP10K+1) expression across all genes in the gene signature. Subsequently, to identify statistically significant gene expression patterns, we computed the mean log_2_(TP10K+1) expression across a background set of 50 genes randomly selected with matching expression levels to those of the genes in the signature iterated 25 times. The gene signature score was defined to be the excess in expression found across the genes in the gene signature compared to the background set.

### Cell-cell Interaction Analysis by Receptor-Ligand Pair Expression

To characterize potential cell-cell interactions, we attempted to identify pairs of cell types where one expresses a receptor gene and the other expresses its cognate ligand. First, we identified known receptor ligand pairs defined in the FANTOM5 receptor ligand database (*142*). Next, we computed a log fold change and p-value for the expression of each gene in each cell type *vs*. profiles from all other cell types to identify genes differentially expressed in each cell type (separately for untreated and treated data). We discarded receptor ligand pairs where either receptor or ligand have a log fold change below 1.5. We then calculated a Receptor-Ligand (RL) score by multiplying the log_2_ fold change of the receptor (in cell type *i*) and ligand (in cell type *j*) to give higher priority to receptor ligand pairs that were highly specific to the respective cell types.

### Consensus Non-Negative Matrix Factorization

We formulated the task of dissecting gene expression programs as a matrix factorization problem where the input gene expression matrix *X*_*n,m*_ is decomposed into two matrices *X*_*n,m*_*=W*_*n,p*_× *H*_*p,m*_*s. t. W, H* > *0*. The solution to this formulation can be identified by solving the following minimization problem:

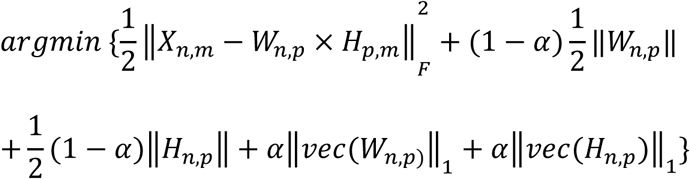

We utilized the non-negative matrix factorization implemented in sklearn to derive the tumor and CAF expression programs. Because the result of NMF optimization can vary between runs based on random seeding, we repeated NMF 50 times per cell type category and computed a set of consensus programs by aggregating results from all 50 runs and computed a stability and reconstruction error. This consensus NMF was performed by making custom updates to the cNMF python package. To determine the optimal number of programs (*p*) for each cell type and condition, we balanced between maximizing stability and minimizing error of the cNMF solution, while ensuring that the resulting programs were as biologically coherent and parsimonious as possible. Each program was annotated utilizing a combination of GSEA (*143*) and comparison to bulk expression signatures.

### Survival Analysis of Bulk RNA-seq Data

Bulk RNA-seq data from two previously published resected primary PDAC cohorts with overall survival annotated were obtained (The Cancer Genome Atlas, *n* = 139; PanCuRx, *n* = 168) (*11, 78*). Patients with metastases or those that received neoadjuvant therapy were excluded from this analysis. Gene expression levels from RNA-seq data was estimated using RSEM (*144*).

To score untreated malignant and fibroblast programs in each tumor, we calculated cNMF-weighted expression scores for each program and normalized the expression scores by calculating z-scores. Notably, for malignant programs we scored the five lineage programs (due to the overlap between cell state and lineage programs and to reduce complexity). For fibroblasts, we scored all four programs.

For the malignant only analysis, we scored each tumor for the five untreated malignant lineage cNMF programs (normalized by z-scores), and identified the top scoring program (primary program) as well as the number of highly-expressed programs defined as expression greater than the mean of the cohort (heterogeneity score, H). Patients with 0 or 1 highly-expressed programs were assigned H = 0, those with 2 highly-expressed programs were assigned H = 1, and those with 3 or more highly-expressed programs were assigned H = 2. We then stratified each tumor into one of 15 groups based on the combination of the primary program (5) and heterogeneity score (3) (**Figure 3E**, row labels). Analogously, for the fibroblast only analysis, we scored each tumor based on primary program (4) and heterogeneity score (2) to stratify each tumor into one of 8 groups (**Figure 3E**, column labels). For the fibroblasts, there was one fewer program than the malignant cells so we scored patients with 0 or 1 high-expressed programs with H = 0 and those with 2 or more high-expressed programs with H = 1.

Next, we partitioned the tumors in the malignant- and fibroblast-only analyses into three risk groups: low, intermediate, high. To this end, we performed preliminary Kaplan-Meier (KM) analyses for overall survival (OS) based on each individual primary program, heterogeneity score, and both (**Figures S9A-F**). We inspected the survival curves to consider any prognosis trends and used this information to assign combinations of primary programs and heterogeneity score values into low, intermediate and high risk groups *pre-hoc* (i.e., prior to the KM analyses of **Figures 3F-G**). This was done separately for the malignant cells (**Figure 3E**, left vertical red color bar) and the fibroblasts (**Figure 3E**, top horizontal blue color bar).

To assign tumors into risk strata based on both malignant and fibroblast program scores and heterogeneity indices, we used the tumor assignment to malignant and fibroblast risk groups above, which defined nine combinations (matrix entries in **Figures 3E, S9G**). We then grouped possible combinations into four risk strata: both low (“low”), both high (“high”), one high and one intermediate (“high-intermediate”), and all other combinations (“low-intermediate”) (color grid inside matrix in **Figure 3E**, purple color legend).

Finally, we performed survival analyses for the risk stratified data based on three malignant risk groups (**Figure 3F**), three fibroblast risk groups (**Figure 3G**), and four combined risk groups (**Figure 3H**). Survival analysis was performed using the Kaplan-Meier estimate and the survival distributions for each patient strata were compared using the log-rank test for equality of survivor functions (Stata/SE 15.1).

### Digital Spatial Profiling (DSP)

We followed published experimental methods (*125*) with modifications as noted below. Briefly, formalin-fixed paraffin-embedded (FFPE) sections (5 µm) of 12 specimens (8 untreated, 4 treated) were prepared by the MGH Histopathology Core on the IRB-approved protocol (2003P001289). Slides were baked at 37°C overnight, deparaffinized, rehydrated, antigen-retrieved in pressure cooker for 20 min at 100°C and low pressure, proteinase-K digestion for 15 min, post-fixed in neutral-buffered formalin for 10 min, hybridized to UV-photocleavable barcode-conjugated RNA *in situ* hybridization probe set (cancer transcriptome atlas/CTA with 1,412 targets or whole transcriptome atlas/WTA) overnight, washed to remove off-target probes, and then counterstained with morphology markers for 2 hours. The morphology markers consisted of: 1:10 SYTO13 (ThermoFisher Scientific, cat. no. 57575), 1:40 anti-panCK-Alexa Fluor 532 (clone AE-1/AE-3; Novus Biologicals, cat. no. NBP2-33200AF532), 1:40 anti-CD45-Alexa Fluor 594 (clone 2B11 + PD7/26; Novus Biologicals, cat. no. NBP2-34528AF594), and 1:100 anti-αSMA-Alexa Fluor 647 (clone 1A4; Novus Biologicals, cat. no. IC1420R). These four morphology markers allowed delineation of the nuclear, epithelial, immune, and fibroblast compartments, respectively. Immunofluorescence images, region of interest (ROI) selection, segmentation into marker-specific areas of interest (AOI), and spatially-indexed barcode cleavage and collection were performed on a GeoMx Digital Spatial Profiling instrument (NanoString) using either the pre-commercial Cancer Transcriptome Atlas (CTA) probe set or a whole transcriptome atlas assay (NanoString). Approximately 8-10 ROIs and 23-25 AOIs were collected per specimen. Library preparation was performed according to the manufacturer’s instructions and involved PCR amplification to add Illumina adapter sequences and unique dual sample indices. Up to 96 AOIs were pooled and sequenced on a NextSeq High Output v2.5 (75 cycles, 2×38 bp; Illumina, cat. no. 20024906).

### Computational Analysis of DSP Data

FASTQ files for DSP were aggregated into count matrices as described previously (*125*). Briefly, deduplicated sequencing counts were calculated based on UMI and molecular target tag sequences. Outlier probes were removed per target when multiple probes were available, and target expression values were calculated as the geometric mean of the remaining probes. Single probe genes were reported as the deduplicated count value. The limit of quantitation (LOQ) was estimated as the geometric mean of the negative control probes plus 2 geometric standard deviations of the negative control probes. Targets were removed that consistently fell below the LOQ, and the datasets were normalized using upper quartile (Q3) normalization.

Statistical analysis was performed using R. For DSP analysis of individual data points, when feasible, linear mixed effect models (*145*) were used to control for multiple sampling within a slide, using Satterthwaite’s approximation (*146*) for degrees of freedom for p-value calculation. When replication was insufficiently powered for mixed effect models, Student’s t-test was used to test associations with subtype classification. All analyses were two-sided and used a significant level of p-value ≤0.05 and were adjusted for multiple testing where appropriate using the false discovery rate (*147*). Programs were scored for each DSP sample within each region of interest using single-sample gene set enrichment analysis (ssGSEA) (*148*). To further align our malignant programs with the conventional classification, comparisons between classical-like and basal-like subtypes were performed after mean-centering the various cNMF program scores within these aggregate categories as described previously.

## DATA AVAILABILITY

Raw data will be available in the controlled access repository Data Use Oversight System (DUOS) at the Broad Institute: https://duos.broadinstitute.org/ under its Data Access Committee. Processed annotated datasets is provided in the Single Cell Portal. The treatment-naïve data is at:https://singlecell.broadinstitute.org/single_cell/study/SCP1089/human-treatment-naive-pdac-snuc-seq and the post-treatment data is at: https://singlecell.broadinstitute.org/single_cell/study/SCP1096/human-treated-pdac-snuc-seq

## CODE AVAILABILITY

All code will be available upon publication in Github at https://github.com/karthikj89/humanpdac.

## ACKNOWLEDGEMENTS

We thank Leslie Gaffney for assistance with preparing figures and Jason Ptacek from IONpath for assistance with MIBI. This work was supported in part by the Ludwig Institute for Cancer Research (A.R.), Klarman Cell Observatory (A.R.), Lustgarten Foundation (T.J.), American Society for Clinical Oncology/Conquer Cancer Foundation Young Investigator Award (W.L.H), UCSF Dean’s Yearlong Fellowship (J.A.G.), and Early Postdoc Mobility Fellowship (no. P2ZHP3 181475) from the Swiss National Science Foundation (D.S.). This study was also conducted with support of the Ontario Institute for Cancer Research (PanCuRx Translational Research Initiative) through funding provided by the Government of Ontario, the Wallace McCain Centre for Pancreatic Cancer supported by the Princess Margaret Cancer Foundation, the Terry Fox Research Institute, the Canadian Cancer Society Research Institute, and the Pancreatic Cancer Canada Foundation. W.L.H. is an Andrew L. Warshaw, M.D. Institute for Pancreatic Cancer Research Fellow. D.S. is a Damon Runyon Cancer Research Fellow (DRQ-03-20). T.J. and A.R. are investigators of the Howard Hughes Medical Institute.

## AUTHOR CONTRIBUTIONS

W.L.H., K.A.J., C.F.C., A.S.L., O.R.R., T.E.J., and A.R. developed the study concept and designed the experiments. K.A.J., W.L.H., J.A.G., H.I.H., T.E.J., and A.R. analyzed the single-nucleus RNA-seq data with computational assistance from P.Y., R.M., O.A., J.G., C.L., C.P., and A.T. and scientific insights from J.S., W.F-P., D.T.T., and A.J.A. W.L.H., G.E., C.R., D.C., C.F.C., and A.S.L. acquired patient-derived tumor specimens with clinical input from M.M.K., A.J.A., T.S.H., J.Y.W., C.R.F., D.T.T., C.D.W., D.P.R, and J.S.L. W.L.H., C.F.C., and A.S.L. curated clinical data. W.L.H., E.D., and N.V.W. developed and optimized the single nucleus RNA-seq method for pancreatic cancer specimens. W.L.H., E.D., N.V.W., J.A.G., H.I.H., D.D., J.W., M.C., L.N., T.D., and D.P. performed single-nucleus RNA-seq experiments, with guidance from O.R.R. S.F. and D.A. performed multiplexed ion beam imaging (MIBI) experiments and D.S., P.Y., K.A.J., W.L.H., J.A.G., and H.I.H. analyzed the data. H.I.H., W.L.H., D.Z., K.F., and R.F. performed the digital spatial profiling (DSP) experiments and J.R., D.Z., W.L.H, and H.I.H. analyzed the data. M.K. and M.M.K. provided histological sections for MIBI and DSP experiments. M.M.K. performed histopathological analyses. W.L.H., K.A.J., J.A.G., H.I.H., T.J., and A.R. wrote the manuscript and all authors reviewed the manuscript.

## MATERIALS & CORRESPONDENCE

Aviv Regev

## SUPPLEMENTAL TABLES

**Supplemental Table 1**. Patient cohort and clinicopathologic data.

**Supplemental Table 2**. Select differentially expressed genes in treated *vs*. untreated tumors, or high *vs*. low residual post-treatment tumors.

**Supplemental Table 3**. Weighted gene lists for cNMF malignant and CAF programs corresponding to Figures 3, S11.

**Supplemental Table 4**. Gene Set Enrichment Analysis results for malignant cell programs ranked by decreasing -log_10_(FDR q-value). Threshold FDR q-value < 0.05.

**Supplemental Table 5**. Gene Set Enrichment Analysis results for CAF programs ranked by decreasing -log_10_(FDR q-value). Threshold FDR q-value < 0.05.

**Supplemental Table 6**. Associations between malignant cell and fibroblast programs and the presence or absence of immune cells or fibroblasts in the same region of interest (ROI) based on the DSP cancer transcriptome atlas data. Statistical tests based on mixed effects model.

**Supplemental Table 7**. Associations between individual immune cell-type genes and epithelial AOI subtype (classical-like *vs*. basal-like) corresponding to Figure 4F. Statistical tests based on Student’s t-test.

## SUPPLEMENTAL FIGURE LEGENDS

**Supplemental Figure 1. Cell Type Composition across PDAC Tumors**. (A) UMAP embeddings of single nucleus profiles (dots) from individual tumors (panels) from untreated (left) and treated (right) patients colored by *post hoc* cell type annotations (color legend). (B,C) Cell type compositions across tumors. Proportion of nuclei (y axis) of each cell type (color legend) in each tumor (x axis) from untreated (left) and treated (right) patients, out of all cells (B) or when considering only non-malignant cells (C).

**Supplemental Figure 2. Inferred CNAs Recover Common Aberrations based on PDAC Genome Studies**. (A) Example inferCNV analysis. Inferred amplifications (red) and deletions (blue) based on expression (color bar) in 100-gene window in each locus (columns) from each cell (rows) labeled by its annotated expression type (color code) in reference cells from matched adjacent normal tissue (top) and cells from the tumor (bottom). (B) Inferred CNA frequencies in our cohort agree with PDAC genome studies. Frequency (y axis) of CNAs on each chromosome arm (x axis) as inferred across the patients in our cohort (light green bars) and from genome analysis of PDAC (dark green bars) and prostate adenocarcinoma (PRAD) (grey bars) from TCGA cohorts.

**Supplemental Figure 3. snRNA-seq Captures Representative Cell Types Distributions Compared to *in situ* Assessment by MIBI**. Proportion of cells (y axis) in each of the four major compartments (color legend) as estimated by snRNA-seq or MIBI (x axis) in each untreated (top; n = 5) or treated (bottom; n = 2) tumor measured by MIBI (2-3 fields of view per slide).

**Supplemental Figure 4. CRT remodels cell type composition across compartments**. Proportions (y axis) of cell subsets (x axis) in untreated (n = 15) *vs*. treated (n = 11) patients out of all non-malignant cells in the tumor (A) or out of all stromal cells only (B). * p < 0.05, Fisher’s exact test.

**Supplemental Figure 5. Treatment Impact of Gene Expression in Different Cell Subsets**. (A-C) Differential expression (log_2_(fold-change), *x* axis) and its significance (-log_10_(adjusted p-value), *y* axis, DESeq2 R package) between treated and untreated tumors (A), CD8^+^ T lymphocytes, (B), macrophages, (C), malignant cells omitting two treated tumors with germline *BRCA2* mutations (PDAC_T_1,2)) or (D), of genes in malignant cells from treated tumors with high (>10%, PDAC_T_5,7,10,11) *vs*. low (PDAC_T_1,2,3,4,6,8,9) residual neoplastic content. Names of selected significant genes are marked.

**Supplemental Figure 6. Bulk-Derived Tumor Subtype Signatures Across Single Nuclei in the PDAC Cohort**. UMAP embeddings of single nucleus profiles (dots) from all tumor nuclei (top two panels) or only malignant cells (bottom two panels) separately for untreated and treated patients colored by expression score (color bar, **Methods**) of signatures derived from the Bailey (*10*), Collisson (*7*), Moffitt (*9*), and COMPASS/PanCuRx (*78*) studies.

**Supplemental Figure 7. cNMF Program Distributions across Malignant Cells in Individual Tumors**. Proportion of cells (*y* axis) assigned with highest scoring program (color legend) in individual tumors (*x* axis) in the untreated (left) and treated (right) groups.

**Supplemental Figure 8. Association between Basal-like and Interferon, TNF-NFkB Programs**. Normalized correlation (color bar) of the gene weights for each cNMF program (rows, columns) in untreated (left) or treated (right) tumors.

**Supplemental Figure 9. Survival Analysis of Bulk RNA-seq PDAC Cohort Based on Malignant Cell and Fibroblast Programs and Heterogeneity Score**. (A-G) Kaplan-Meier survival analyses of PDAC cohort (n = 307) from TCGA (*11*) and PanCuRx (*78*), stratified by primary malignant program (A), malignant heterogeneity score (B), combined primary malignant program and heterogeneity score (C), primary fibroblast program (D), fibroblast heterogeneity score (E), combined primary fibroblast program and heterogeneity score (F), and combined primary malignant program and heterogeneity score with combined primary fibroblast program and heterogeneity score (G). Survival distributions for each patient strata were compared using the log-rank test.

**Supplemental Figure 10. Previous CAF Subset Signatures across Single Fibroblast Profiles in the PDAC Cohort**. UMAP embeddings of single nucleus profiles (dots) from fibroblast nuclei from untreated (top) and treated (bottom) patients colored by expression score (color bar, **Methods**) of three signatures previously reported by Tuveson and colleagues (*51*).

**Supplemental Figure 11. Differences in Fibroblast Gene Expression, Composition and Programs in Treated Tumors**. (A) Cell intrinsic expression differences in fibroblasts from CRT-treated tumors. Left: differential expression (log_2_(fold-change), x axis) and its significance (-log_10_(adjusted p-value), *y* axis, DESeq2 R package) of genes in fibroblasts between treated and untreated tumors. Names of selected significant genes are marked. Right: GSEA (*143, 149, 150*) terms (y axis) ranked by increasing significance (-log_10_(FDR q-value)) of enrichment in treated tumors. (B) cNMF expression program dictionary in fibroblasts from untreated and treated tumors. UMAP embeddings of single nucleus profiles (dots) from untreated (top) and treated (bottom) tumors, colored by patient (left panel) or by the score derived for each cell-cNMF program pair (color bar, **Methods**). (C) Normalized correlation (color bar) of the gene weights for each cNMF program (rows, columns) in untreated (top) or treated (bottom) tumors. (D) Higher proportion of myofibroblast and neuromuscular programs in CRT treated tumors. Proportion of fibroblasts primarily expressing each fibroblast cNMF program within untreated (left) and treated (right) tumors, in aggregate across all tumors (top) or in individual tumors (bottom, x axis).

**Supplemental Figure 12. Assessing PDAC Programs by Digital Spatial Profiling**. (A) Experimental workflow for digital spatial profiling on the GeoMx platform (NanoString). (B) Spatial resolution of cell types across ROIs and AOIs. Expression (z-score of normalized counts across AOIs; purple/yellow color bar) of signature genes (rows) from diverse cell types (color legend (4) and left bar) across AOIs (columns, color legend and horizontal bar (3)) profiled by 1,412-gene cancer transcriptome atlas or CTA, capturing epithelial (green), fibroblasts (blue) and immune (red) cells, from ROIs characterized by presence or absence of immune (color legend and horizontal bar (1)) and fibroblast (color legend and horizontal bar (2)) infiltration. Both columns and rows are clustered by unsupervised hierarchical clustering. (C) Box-plots comparing mean normalized gene expression by cluster for basal-like *vs*. classical-like epithelial AOIs. * p < 0.05, Student’s t-test. (D) Coverage of PDAC snRNA-seq programs by CTA and WTA digital spatial profiling. Number of genes (y axis) from each untreated malignant cell program (x axis) captured by CTA only (white), WTA only (black), or both (grey). (E) Impact of gene panel on program scores. Spearman correlation coefficient (ρ) between the scores for different untreated malignant programs (x axis) obtained with WTA using the full gene panel *vs*. the gene subset shared with the CTA assay.

