## Supplemental Figures 1-12 for "Single-nucleus and spatial transcriptomics of archival pancreatic cancer reveals multi-compartment reprogramming after neoadjuvant treatment"

### Supplemental Figure 1

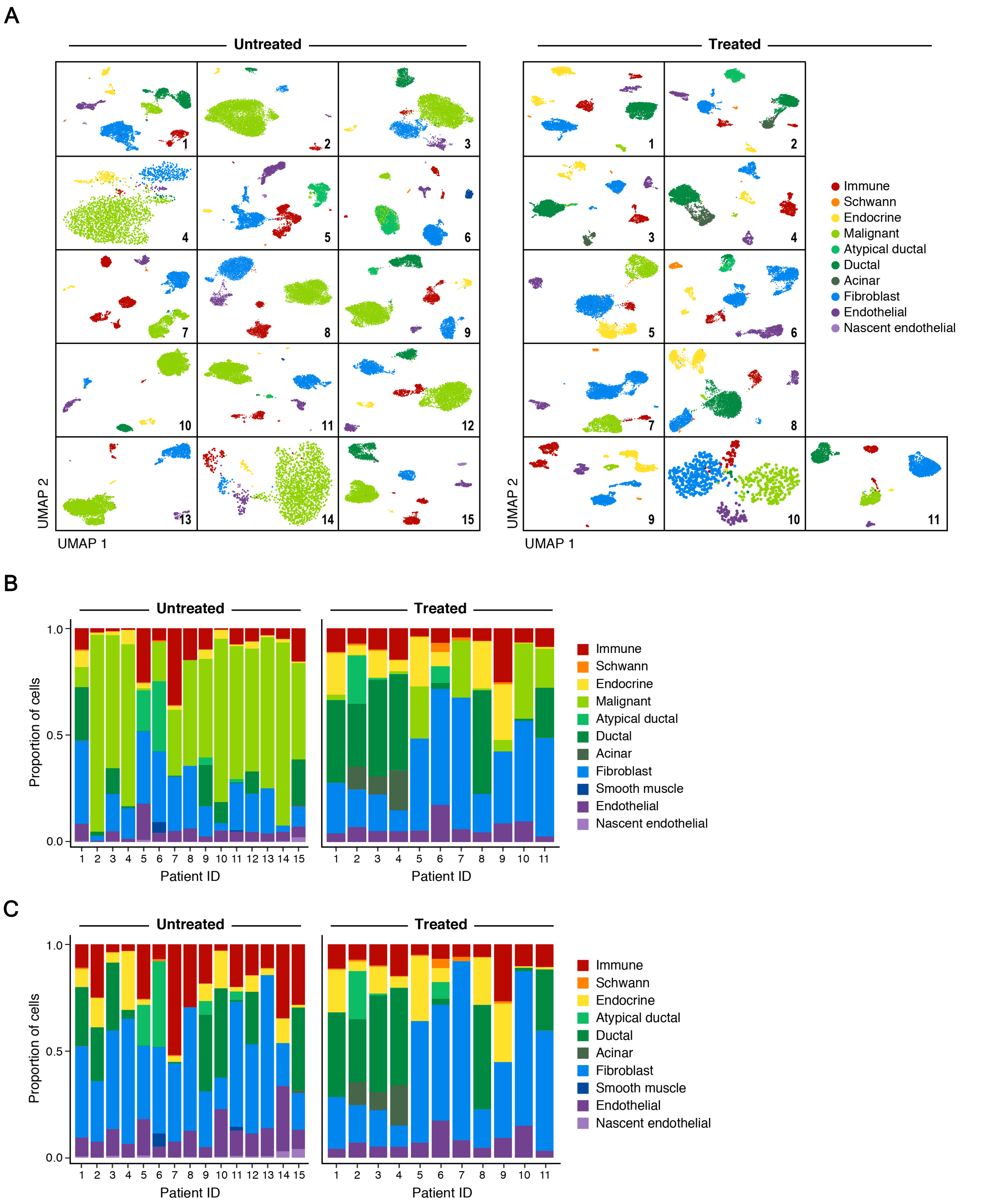

### Supplemental Figure 2

A

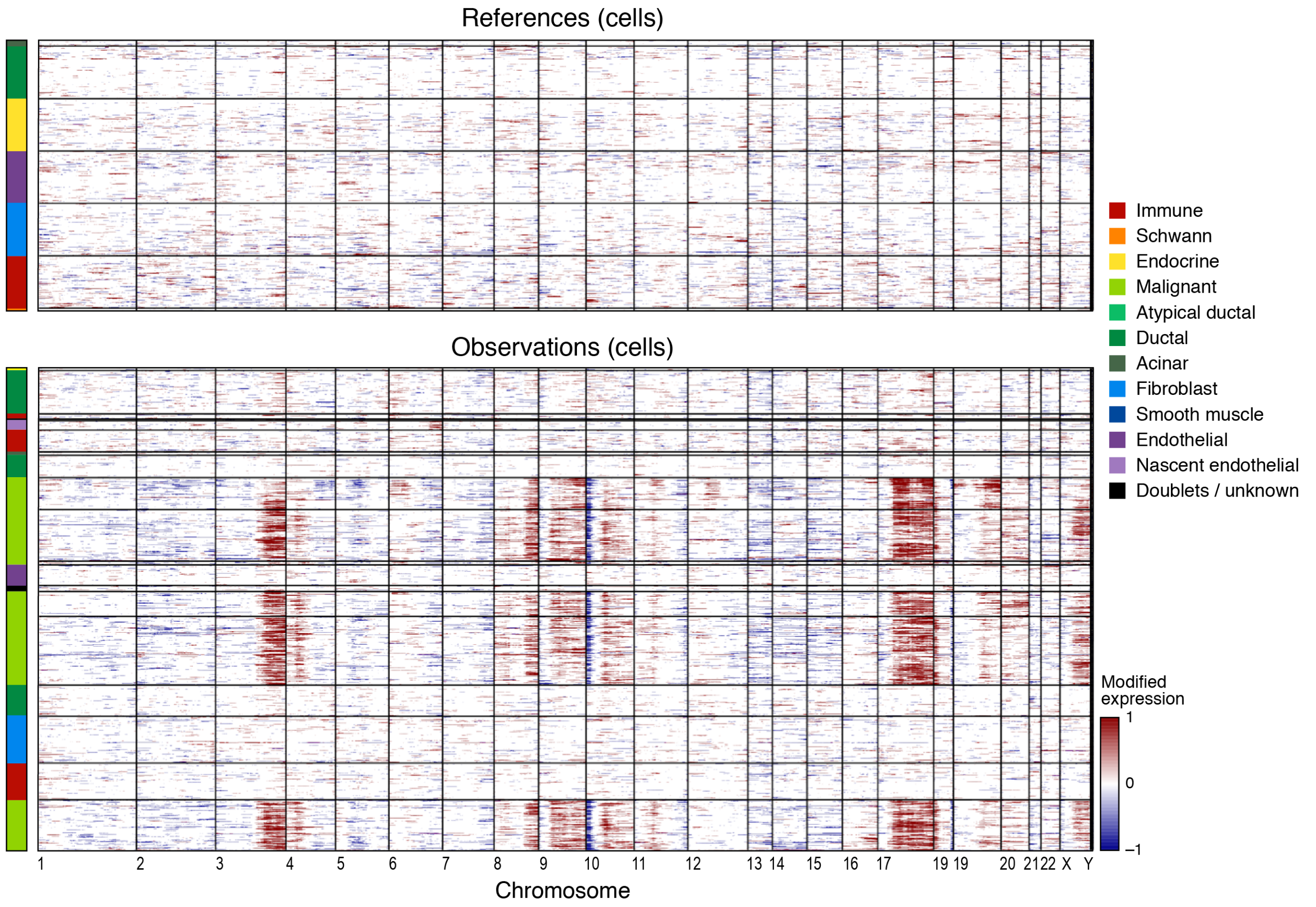


B

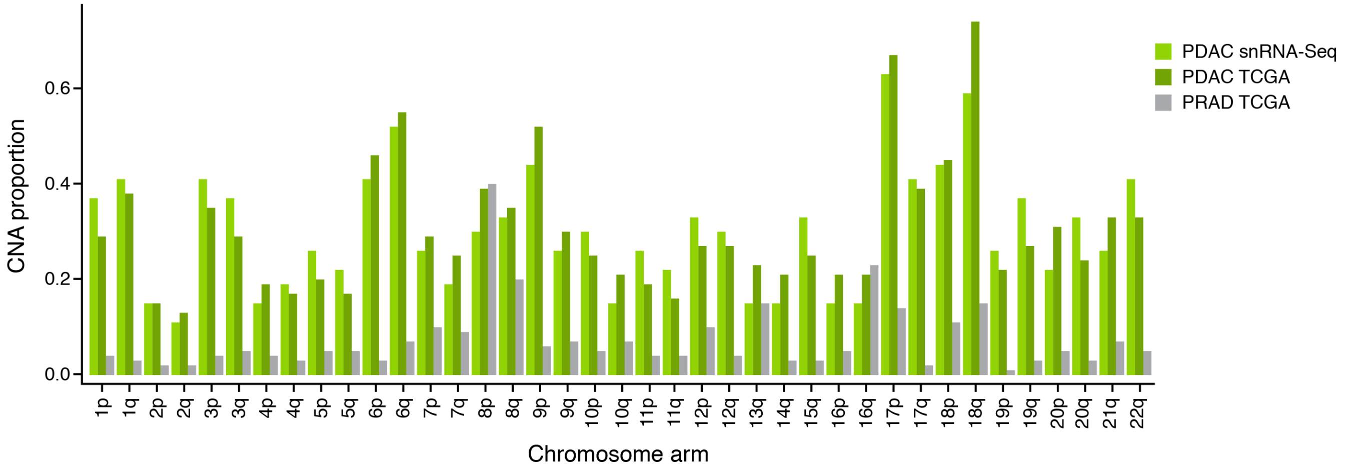

### Supplemental Figure 3

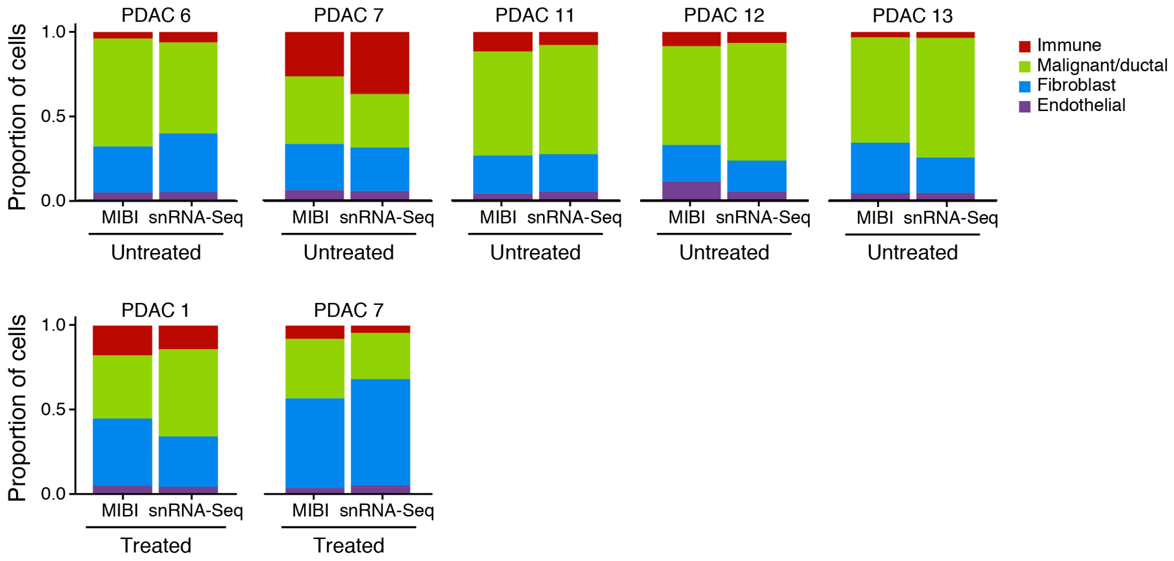

### Supplemental Figure 4

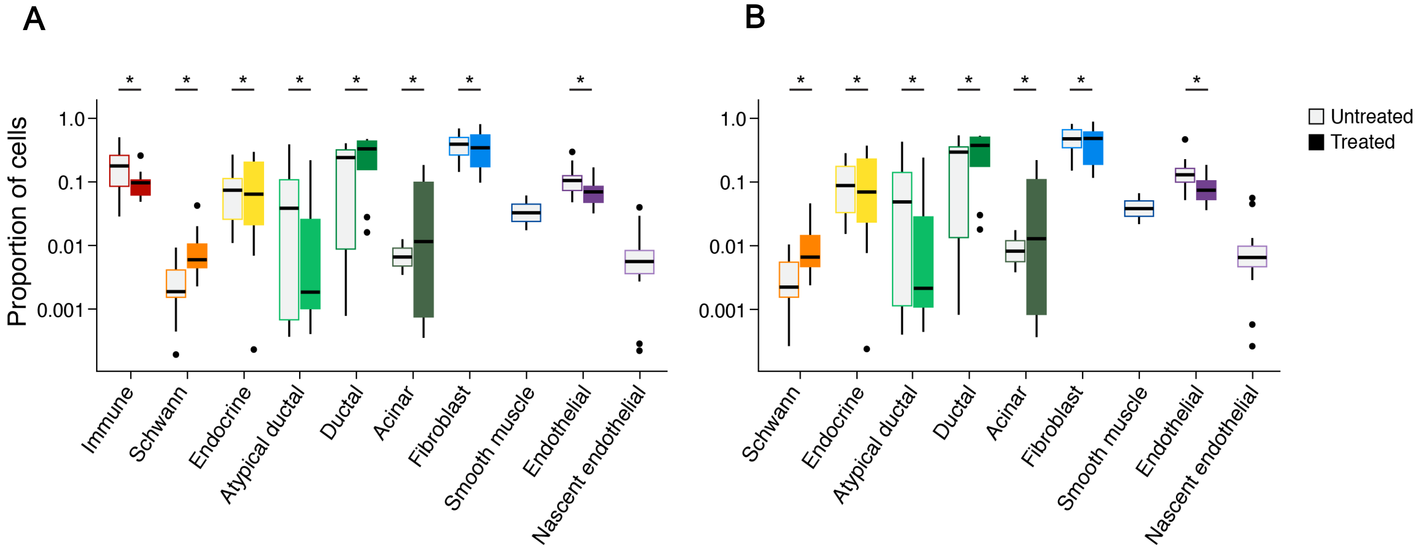

### Supplemental Figure 5

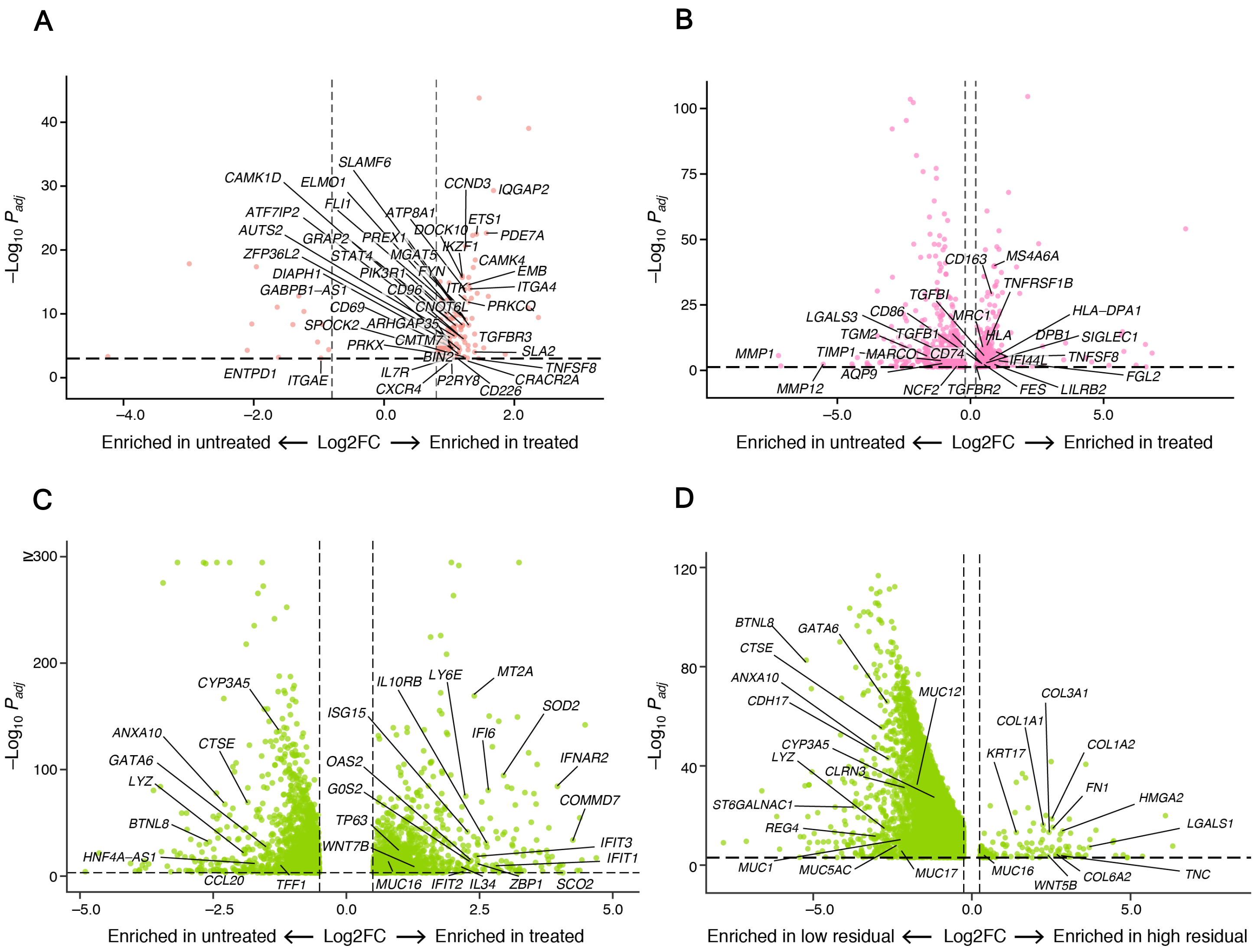

### Supplemental Figure 6

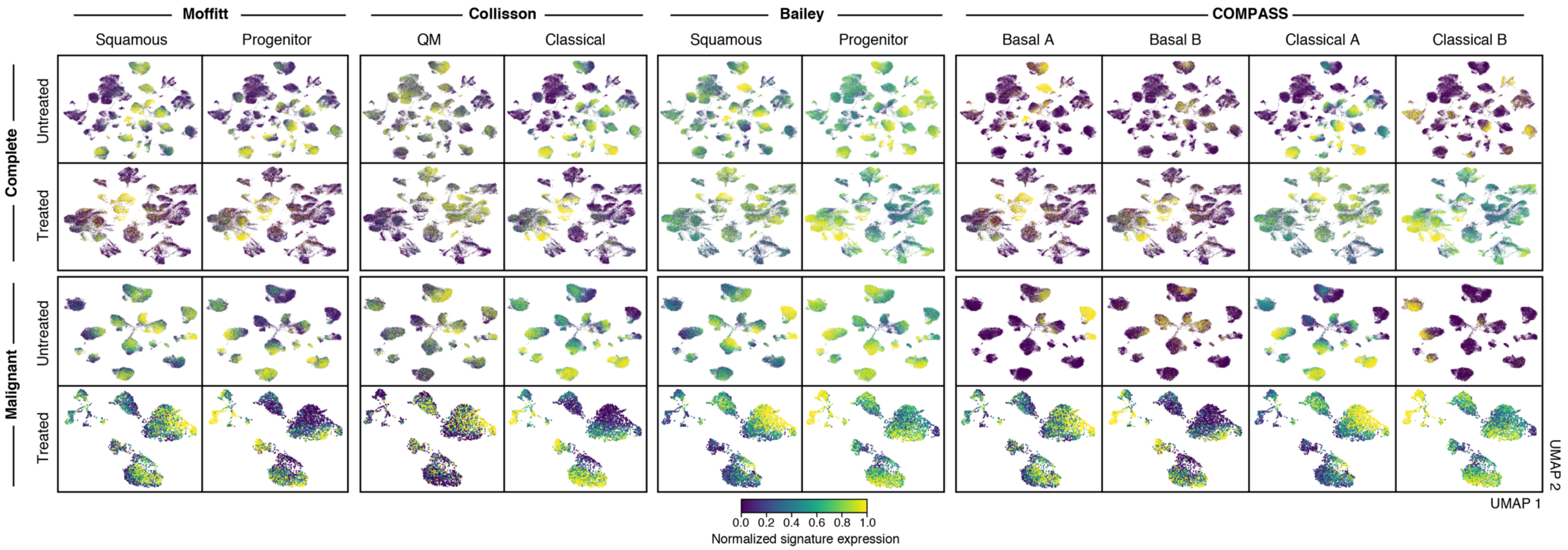

### Supplemental Figure 7

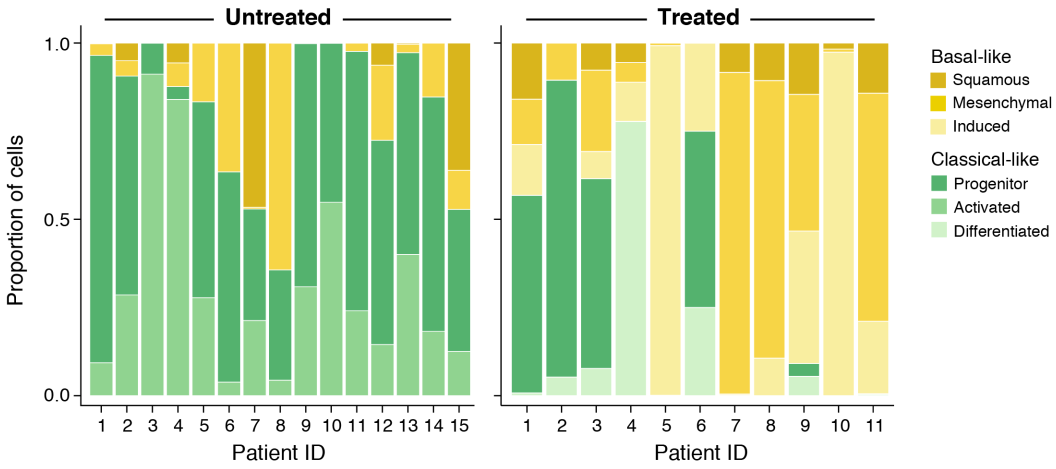

### Supplemental Figure 8

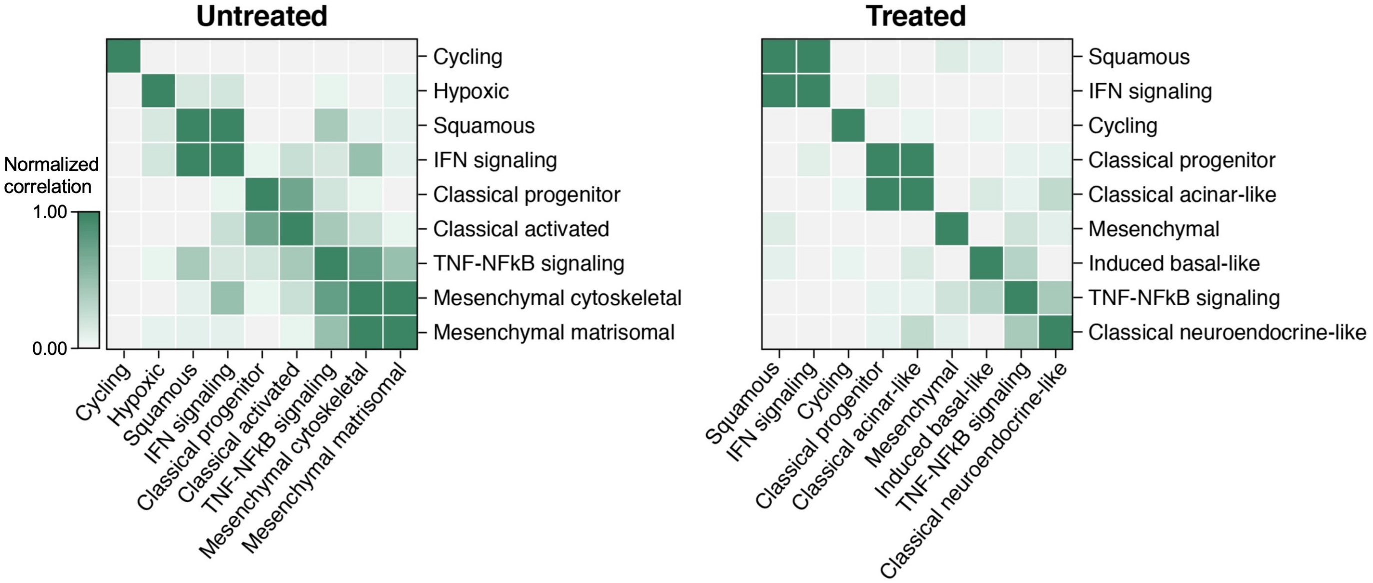

### Supplemental Figure 9

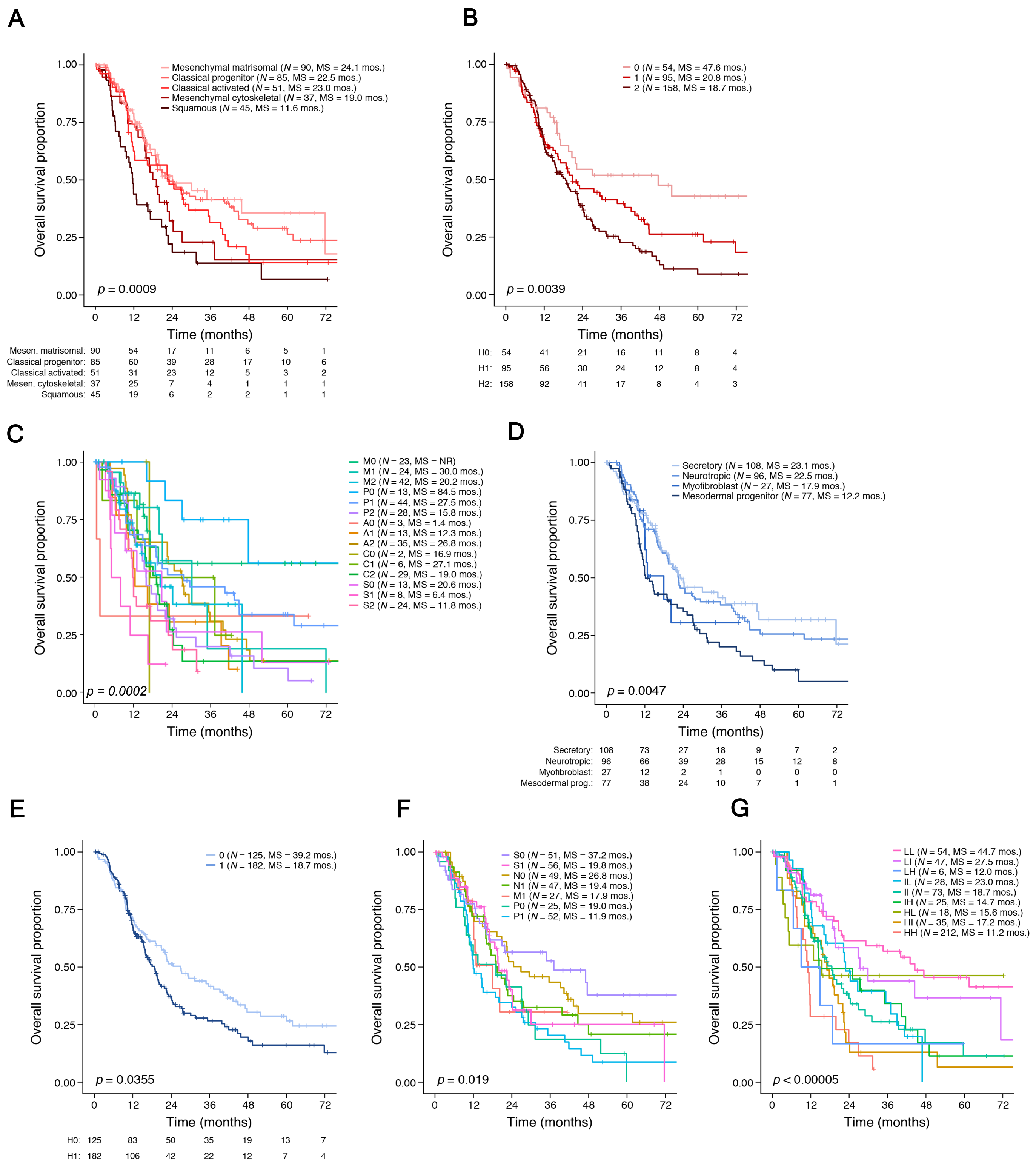

### Supplemental Figure 10

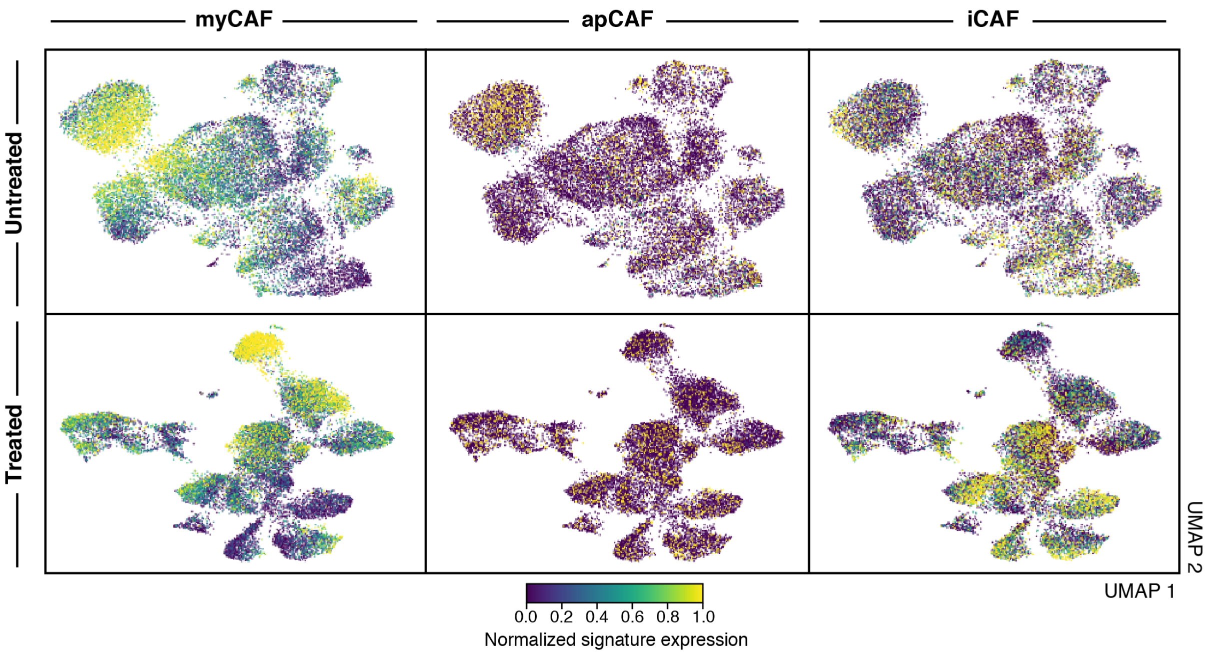

### Supplemental Figure 11

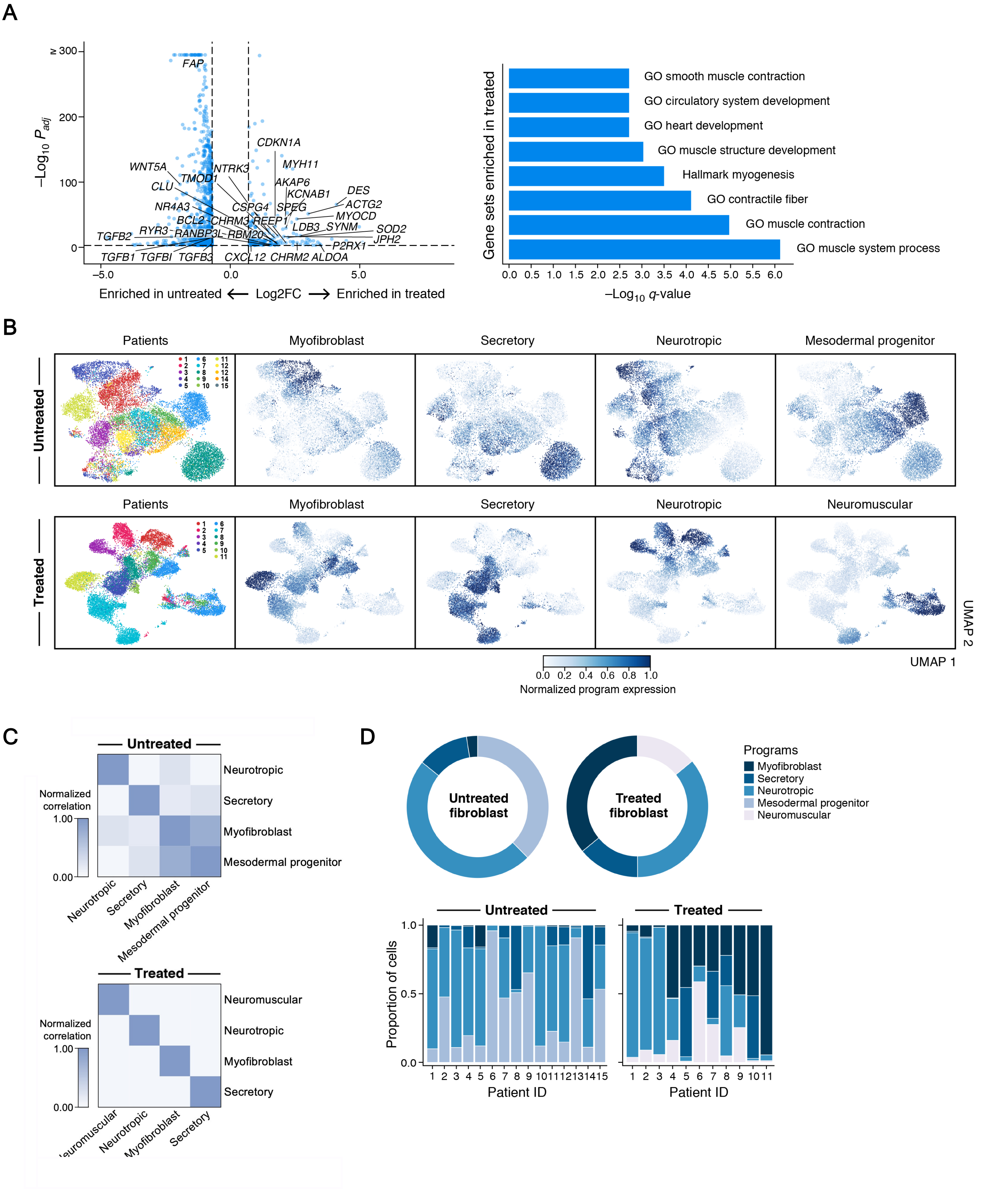

### Supplemental Figure 12

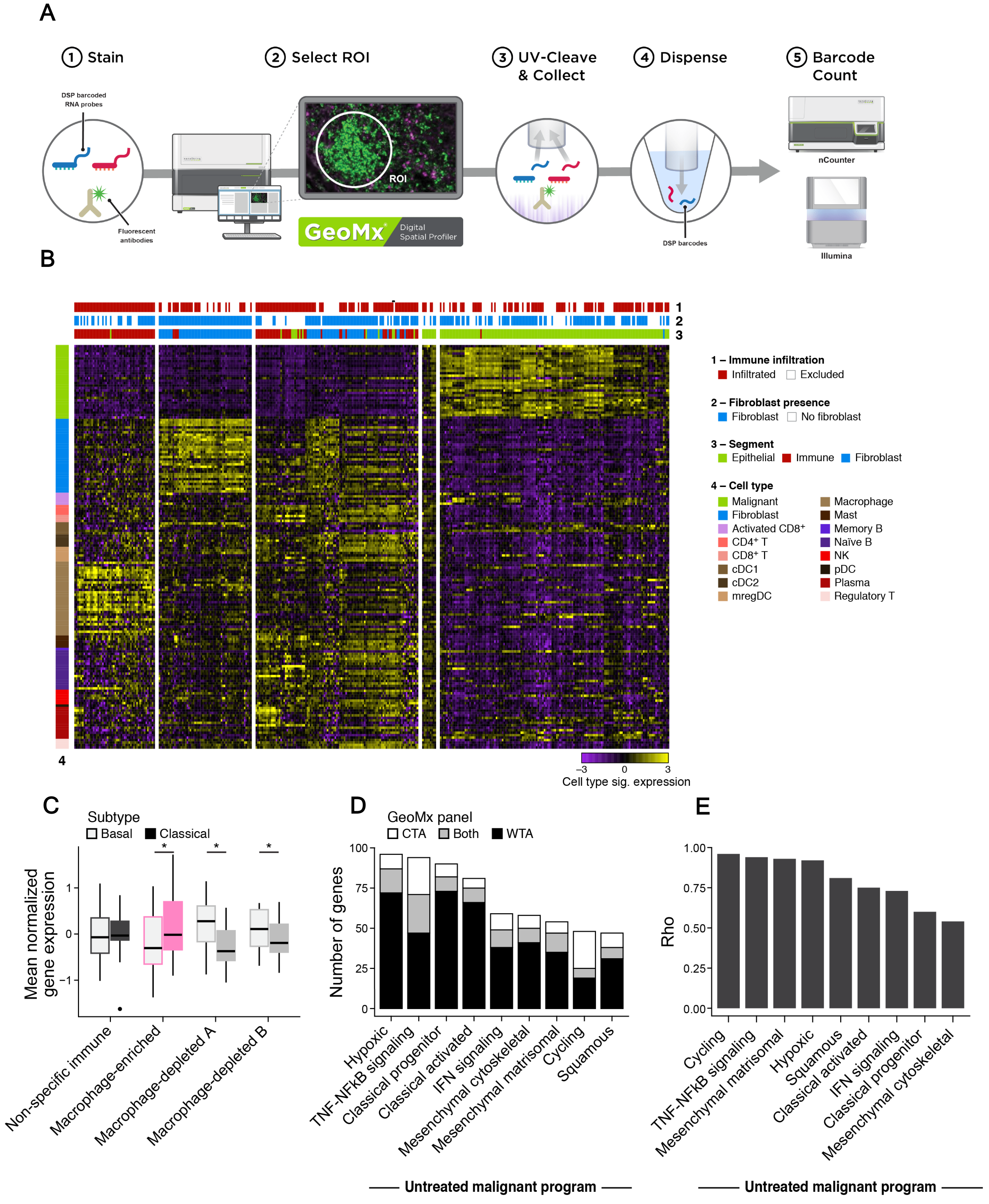
