## Supplemental Tables 1,4,5 for "Single-nucleus and spatial transcriptomics of archival pancreatic cancer reveals multi-compartment reprogramming after neoadjuvant treatment"

### Supplemental Table 1

| ID | Age (decade) | Sex | Stage/Grade | Margin | Histology | Neoadjuvant | Adjuvant | Status Last FUP | PFS (d) | OS (d) | Storage (d) | 10x Chemistry |
|---|---|---|---|---|---|---|---|---|---|---|---|---|
| **Untreated (U)** | | | | | | | | | | | | |
| PDAC_U_1 | 20s | F | T3N1M0/g2-3 | R1 | | None | FOLFIRINOX/Gem/RT | MET | 70 | 892 | 638 | v3 |
| PDAC_U_2 | 60s | F | T3N2M0/g2 | R1 | AS | None | FOLFIRINOX/RT | LR/DWD | 78 | 297 | 17 | v3 |
| PDAC_U_3 | 60s | F | T3N0M0/gX | R0 | | None | Gem/cape/RT | MET | 578 | 649 | 341 | v3 |
| PDAC_U_4 | 60s | M | T3N1M0/g3 | R0 | | None | Gem/cape/RT | LR/DWD | 267 | 538 | 634 | v2 |
| PDAC_U_5 | 60s | M | T3N1M0/g3 | R1 | | None | Gem/cape/RT | NED | 282 | 282 | 661 | v3 |
| PDAC_U_6 | 60s | M | T2N2M0/g2 | R0 | | None | Gem/cape | NED | 151 | 151 | 220 | v3 |
| PDAC_U_7 | 70s | M | T3N2M0/g2 | R0 | AS | None | None | MET/DWD | 208 | 563 | 384 | v3 |
| PDAC_U_8 | 70s | M | T2N1M0/g2 | R0 | | None | Gem/abraxane | MET | 35 | 186 | 62 | v3 |
| PDAC_U_9 | 70s | M | T3N0M0/g2-3 | R1 | | None | None | DWOD | 494 | 494 | 472 | v3 |
| PDAC_U_10 | 70s | F | T2N1M0/g2 | R1 | | None | Gem/cape/RT | NED | 644 | 644 | 393 | v3 |
| PDAC_U_11 | 70s | F | T2N1M0/g2 | R1 | | None | FOLFIRINOX/RT | DWOD | 172 | 172 | 64 | v3 |
| PDAC_U_12 | 70s | M | T2N0M0/g2-3 | R1 | | None | FOLFIRINOX | LR | 329 | 555 | 135 | v2 |
| PDAC_U_13 | 70s | M | T3N0M0/g2 | R1 | | None | Gem/abraxane | NED | 224 | 224 | 106 | v3 |
| PDAC_U_14 | 80s | F | T2N0M0/g2 | R0 | | None | Gem | NED | 225 | 225 | 38 | v2 |
| PDAC_U_15 | 80s | M | T3N2M0/g3 | R1 | AS | None | None | NED | 35 | 35 | 148 | v3 |
| **Treated (T)** | | | | | | | | | | | | |
| PDAC_T_1 | 30s | F | ypT2N0M0/g2 | R0 | BRCA2 germ | FOLFIRINOX/RT | Olaparib | NED | 301 | 301 | 69 | v3 |
| PDAC_T_2 | 50s | M | ypT1cN0M0/g3 | R1 | BRCA2 germ | FOLFIRINOX/RT | FOLFIRINOX | MET | 285 | 285 | 170 | v3 |
| PDAC_T _3 | 50s | F | ypT3N0M0/g2 | R0 | | FOLFIRINOX/RT | None | NED | 1255 | 1255 | 1121 | v3 |
| PDAC_T _4 | 60s | F | ypT3N0M0/g2 | R0 | | FOLFIRINOX/RT | None | NED | 1342 | 1342 | 1142 | v3 |
| PDAC_T _5 | 60s | F | ypT2N0M0/gX | R1 | | FOLFIRINOX/RT | Gem/abraxane | LR | 187 | 433 | 104 | v2 |
| PDAC_T _6 | 60s | F | ypT1aN1M0/gX | R0 | | FOLFIRINOX/RT/nivo | Nivolumab | NED | 264 | 264 | 58 | v3 |
| PDAC_T_7 | 60s | M | ypT2N2M0/g3 | R0 | AS | FOLFIRINOX/RT | None | MET | 34 | 34 | 112 | v2 |
| PDAC_T_8 | 60s | M | ypT1cN0M0/g2 | R0 | | FOLFIRINOX/RT | Gem/abraxane | MET | 117 | 345 | 56 | v2 |
| PDAC_T_9 | 70s | M | ypT3N0M0/g2 | R0 | | FOLFIRINOX/RT/los | None | LR/DWD | 258 | 362 | 905 | v3 |
| PDAC_T_10 | 70s | M | ypT3N0M0/g2 | R0 | | RT/cape | Gemcitabine | Unknown | | | 2389 | v2 |
| PDAC_T_11 | 70s | F | ypT2N1M0/g3 | R0 | | RT | Gemcitabine | MET | 110 | 185 | 365 | v3 |

Abbreviations: AS, adenosquamous; MET, distant metastases; LR, local recurrence; DWD, dead with disease; DWOD, dead without evidence of disease; NED, no evidence of disease; Gem, gemcitabine; Cape, capecitabine; Los, losartan; RT, radiotherapy

### Supplemental Table 4

| Topic | Gene ontology term | -Log(q-value) |
|---|---|---|
| **Untreated** | | |
| Squamous | Charafe breast cancer luminal vs basal dn | 18.8 |
| | Huper breast basal bs luminal up | 16.4 |
| | Hollern squamous breast tumor | 15.9 |
| | GO epidermis development | 12.2 |
| | GO epithelial cell differentiation | 12.0 |
| | GO epithelium development | 11.2 |
| | GO skin development | 11.1 |
| | GO keratinocyte development | 10.4 |
| | GO epidermal cell differentiation | 10.3 |
| | Smid breast cancer basal up | 10.1 |
| Mesenchymal cytoskeletal | Module 47 (ECM and collagens) | 23.7 |
| | Hallmark epithelial mesenchymal transition | 22.4 |
| | Anastassiou multicancer invasiveness signature | 22.1 |
| | Naba matrisome | 17.2 |
| | GO extracellular matrix | 15.9 |
| | GO locomotion | 15.5 |
| | GO supramolecular fiber organization | 15.5 |
| | GO actin filament based process | 14.9 |
| | GO actin binding | 14.2 |
| | GO cell motility | 13.2 |
| Mesenchymal matrisomal | Hallmark epithelial mesenchymal transition | 65.6 |
| | Anastassiou multicancer invasiveness signature | 64.3 |
| | Module 47 (ECM and collagens) | 59.6 |
| | GO extracellular structure organization | 47.0 |
| | GO collagen containing extracellular matrix | 46.7 |
| | GO extracellular matrix | 42.5 |
| | NABA matrisome | 41.7 |
| | NABA core matrisome | 38.9 |
| | GO extracellular matrix structural constituent | 37.2 |
| | Reactome extracellular matrix organization | 36.0 |
| Classical progenitor | Vecchi gastric cancer advanced vs early dn | 19.4 |
| | PRDM6 target genes | 17.5 |
| | DLX2 target genes | 14.0 |
| | SMID breast cancer basal dn | 12.8 |
| | HNF1A target genes | 12.3 |
| | GO plasma membrane region | 10.3 |
| | GATA1 03 | 8.5 |
| | HNF1 01 | 7.5 |
| | Charafe breast cancer luminal vs mesenchymal up | 6.9 |
| | GO ion transport | 4.6 |
| Classical activated | Dodd nasopharyngeal carcinoma up | 13.2 |
| | GO cell leading edge | 5.8 |
| | GO apical part of cell | 4.5 |
| | GO secretory vesicle | 4.2 |
| | GO cell projection membrane | 3.9 |
| | GO secretion | 3.9 |
| | GO cytoskeletal protein binding | 3.9 |
| | HNF1A target genes | 3.6 |
| | GO actin binding | 3.0 |
| | Smid breast cancer basal dn | 2.9 |
| Cycling | Fischer dream targets | 285.9 |
| | GO cell cycle | 139.1 |
| | Benporath cycling genes | 126.6 |
| | GO mitotic cell cycle | 123.3 |
| | GO cell cycle process | 122.3 |
| | Reactome cell cycle | 105.1 |
| Hypoxic | Hallmark hypoxia | 50.5 |
| | Mense hypoxia up | 49.1 |
| | Nakamura tumor zone | 47.7 |
| | Elvidge hypoxia up | 46.9 |
| | Winter hypoxia metagene | 42.6 |
| | Hallmark glycolysis | 21.8 |
| TNF-NFkB signaling | Zhang response to IKK inhibitor and TNF up | 43.0 |
| | Hallmark TNFA signaling via NFkB | 32.0 |
| | Phong TNF response not via p38 | 27.5 |
| | Phong TNF targets up | 22.8 |
| | Charafe breast cancer luminal vs basal dn | 21.4 |
| | Phong TNF response via p38 partial | 20.7 |
| | Hinata NFkB targets keratinocyte up | 18.7 |
| | GO epithelium development | 17.9 |
| | Tian TNF signaling via NFkB | 15.6 |

| | | | |
|---|---|---|---|
| | | GO cytokine mediated signaling pathway | 14.9 |
| | IFN signaling | Hallmark interferon gamma response | 25.7 |
| | | Sana response to IFNG up | 20.9 |
| | | Hecker IFNB1 targets | 19.3 |
| | | Hallmark interferon alpha response | 17.5 |
| | | Browne interferon responsive genes | 16.9 |
| | | Zhang interferon response | 13.8 |
| | | Reactome interferon alpha beta signaling | 13.2 |
| | | Reactome interferon signaling | 12.3 |
| | | GO response to type I interferon | 11.5 |
| | | Einav interferon signature in cancer | 11.3 |
| **Treated** | | | |
| | Squamous | Koinuma targets of SMAD2 or SMAD3 | 11.5 |
| | | Pece mammary stem cell up | 11.5 |
| | | Wu cell migration | 8.8 |
| | | Enk UV response keratinocyte up | 8.4 |
| | | Huper breast basal vs luminal up | 5.6 |
| | | GO epithelium development | 5.3 |
| | | GO epithelial cell differentiation | 4.2 |
| | Mesenchymal | Charafe breast cancer luminal vs basal dn | 11.9 |
| | | Vecchi gastric cancer advanced vs early up | 11.4 |
| | | Charafe breast cancer luminal vs mesenchymal dn | 10.9 |
| | | Nakamura tumor zone peripheral vs central up | 8.6 |
| | | Hallmark epithelial mesenchymal transition | 8.6 |
| | | Reactome extracellular matrix organization | 8.3 |
| | | Module 47 (ECM and collagens) | 8.0 |
| | | Wu cell migration | 8.0 |
| | | GO extracellular structure organization | 7.1 |
| | | GO extracellular matrix | 6.0 |
| | Induced basal-like | Dodd nasopharyngeal carcinoma up | 18.5 |
| | | Wu cell migration | 7.8 |
| | | Charafe breast cancer basal vs mesenchymal up | 7.3 |
| | | Bosco epithelial differentiation module | 5.8 |
| | | HNF1A target genes | 5.3 |
| | | Reactome developmental biology | 5.0 |
| | | Smid breast cancer basal up | 4.6 |
| | | Enk UV response keratinocyte up | 4.4 |
| | | GO response to drug | 3.8 |
| | | GO response to organic cyclic compound | 3.8 |
| | Classical progenitor | Vecchi gastric cancer advanced vs early dn | 10.9 |
| | | GO plasma membrane region | 5.9 |
| | | GO ion transport | 5.9 |
| | | HNF1 01 | 4.2 |
| | | Dodd nasopharyngeal carcinoma up | 3.3 |
| | | Smid breast cancer basal dn | 2.9 |
| | | DLX2 target genes | 2.4 |
| | | HNF1A target genes | 2.3 |
| | | Smid breast cancer luminal b up | 2.3 |
| | | GO apical part of cell | 2.2 |
| | Classical acinar-like | GNF2 SPINK1 | 28.1 |
| | | GNF2 SERPINI2 | 19.3 |
| | | GO digestion | 8.7 |
| | | Smid breast cancer basal dn | 5.0 |
| | | Reactome digestion of dietary lipid | 4.3 |
| | | GO digestive system process | 4.1 |
| | | GO proteolysis | 3.8 |
| | | GATA6 01 | 3.7 |
| | | GO peptidase activity | 3.1 |
| | | Reactome digestion | 2.4 |
| | Classical neuroendocrine-like | GO synapse | 13.8 |
| | | Reactome neuronal system | 11.2 |
| | | GO synaptic signaling | 9.1 |
| | | GO neuron differentiation | 7.4 |
| | | GO vesicle mediated transport in synapse | 7.3 |
| | | GO neuron development | 6.7 |
| | | Reactome regulation of insulin secretion | 5.7 |
| | | GO peptide hormone secretion | 3.1 |
| | | Hallmark pancreas beta cells | 3.1 |
| | | GO regulation of insulin secretion | 2.9 |
| | Cycling | Fischer dream targets | 220.7 |
| | | GO cell cycle | 111.6 |
| | | Benporath cycling genes | 110.4 |
| | | GO mitotic cell cycle | 101.8 |

| | | |
|---|---|---|
| | GO cell cycle process | 94.1 |
| | Reactome cell cycle | 85.0 |
| | GO cell division | 73.1 |
| | Hallmark E2F targets | 72.9 |
| TNF-NFkB signaling | Zwang class 3 transiently induced by EGF | 22.3 |
| | Charafe breast cancer luminal vs basal dn | 13.9 |
| | GO locomotion | 12.5 |
| | Phong TNF response not via p38 | 12.5 |
| | GO cell motility | 10.9 |
| | Zhang response to IKK inhibitor and TNF up | 9.0 |
| | Hallmark TNFA signaling via NFkB | 8.6 |
| | GO regulation of epithelial cell migration | 5.1 |
| | GO taxis | 5.0 |
| | Smid breast cancer basal up | 4.9 |
| IFN signaling | Reactome interferon signaling | 7.3 |
| | Hallmark interferon alpha response | 6.9 |
| | Reactome innate immune system | 6.6 |
| | Reactome interferon alpha beta signaling | 5.4 |
| | Hallmark interferon gamma response | 5.3 |
| | Reactome antigen processing cross presentation | 4.5 |
| | Reactome AP folding assembly and peptide loading of class I MHC | 3.6 |
| | Reactome cytokine signaling in immune system | 3.6 |
| | Reactome interferon signaling | 7.3 |
| | Hallmark interferon alpha response | 6.9 |

### Supplemental Table 5

| Topic | Gene ontology term | -Log(q-value) |
| --- | --- | --- |
| **Untreated** | | |
| Myofibroblast | Hallmark epithelial mesenchymal transition | 35.2 |
| | GO extracellular matrix | 26.5 |
| | GO biological adhesion | 24.4 |
| | GO collagen containing extracellular matrix | 24.3 |
| | GO locomotion | 24.0 |
| | Naba matrisome | 20.7 |
| | GO cell motility | 20.4 |
| | GO actin cytoskeleton | 18.1 |
| Secretory | Reactome eukaryotic translation elongation | 86.4 |
| | Kegg ribosome | 81.1 |
| | GO SRP dependent cotranslational protein targeting to membrane | 78.0 |
| | GO establishment of protein localization to endoplasmic reticulum | 74.9 |
| | GO translational initiation | 63.9 |
| | GO protein targeting to membrane | 62.1 |
| | GO secretory granule | 17.5 |
| | GO secretory vesicle | 16.1 |
| | GO exocytosis | 14.7 |
| | GO cytokine mediated signaling pathway | 11.7 |
| Neurotropic | GO neurogenesis | 13.9 |
| | GO neuron differentiation | 13.4 |
| | GO neuron development | 12.1 |
| | GO synapse | 9.6 |
| | GO synapse organization | 9.4 |
| | GO postsynapse | 9.1 |
| Mesodermal progenitor | PRDM6 target genes | 24.1 |
| | Lim mammary stem cell up | 17.3 |
| | GO embryo development | 16.7 |
| | Boquest stem cell up | 15.2 |
| | GO skeletal system development | 14.4 |
| | GO circulatory system development | 12.7 |
| | GO cardiovascular system development | 11.1 |
| | GO cartilage development | 10.0 |
| | GO connective tissue development | 9.4 |
| | GO mesenchyme development | 7.4 |
| **Treated** | | |
| Myofibroblast | Hallmark epithelial mesenchymal transition | 24.3 |
| | GO biological adhesion | 18.8 |
| | GO locomotion | 18.8 |
| | GO cell motility | 18.6 |
| | GO extracellular structure organization | 16.9 |
| | Naba matrisome | 16.2 |
| | GO cell adhesion molecule binding | 15.0 |
| | GO extracellular matrix | 14.5 |
| | GO collagen containing extracellular matrix | 13.9 |
| | GO response to wounding | 3.4 |
| Secretory | Reactome eukaryotic translation elongation | 125.7 |
| | GO cotranslational protein targeting to membrane | 115.8 |
| | GO establishment of protein localization to ER | 111 |
| | GO protein localization to membrane | 92.6 |
| | GO peptide biosynthetic process | 65.4 |
| | GO secretory vesicle | 12.4 |
| | GO exocytosis | 12.2 |
| | GO secretory granule | 12.1 |
| | GO response to cytokine | 10.6 |
| | GO secretion | 7.7 |
| Neurotropic | GO neurogenesis | 14.9 |
| | GO neuron differentiation | 12.9 |
| | GO neuron development | 10.1 |
| | GO cell projection organization | 9.4 |
| | GO postsynapse | 7.6 |
| Neuromuscular | GO actin filament based process | 16.5 |
| | GO muscle system process | 16.1 |
| | GO muscle structure development | 14.4 |
| | GO contractile fiber | 13.5 |
| | GO muscle contraction | 13.4 |
| | GO I band | 12.1 |
| | GO action potential | 11.1 |
| | GO regulation of membrane potential | 11.0 |
| | GO synapse | 10.4 |
| | GO muscle cell differentiation | 10.1 |
